# Reduced chromatin accessibility underlies gene expression differences in homologous chromosome arms of hexaploid wheat and diploid *Aegilops tauschii*

**DOI:** 10.1101/571133

**Authors:** Fu-Hao Lu, Neil McKenzie, Laura-Jayne Gardiner, Ming-Cheng Luo, Anthony Hall, Michael W Bevan

## Abstract

Polyploidy has been centrally important in driving the evolution of plants, and leads to alterations in gene expression that are thought to underlie the emergence of new traits. Despite the common occurrence of these global patterns of altered gene expression in polyploids, the mechanisms involved are not well understood. Using a precise framework of highly conserved syntenic genes on hexaploid wheat chromosome 3DL and its progenitor 3L chromosome arm of diploid *Aegilops tauschii*, we show that 70% of these genes exhibited proportionally reduced gene expression, in which expression in the hexaploid context of the 3DL genes was approximately 40% of the levels observed in diploid *Ae. tauschii.* Many genes showing elevated expression during later stages of grain development in wheat compared to *Ae. tauschii.* Gene sequence and methylation differences accounted for only a few cases of differences in gene expression. In contrast, large scale patterns of reduced chromatin accessibility of genes in the hexaploid chromosome arm compared to its diploid progenitor were correlated with observed overall reduction in gene expression and differential gene expression. Therefore, that an overall reduction in accessible chromatin underlies the major differences in gene expression that results from polyploidization.

## Introduction

Polyploidy arises from the duplication of genomes or the fusion of genomes from different species and has occurred frequently in the lineages of many organisms, from fish to flowering plants (Otto 2007). The ancestral flowering plant lineage underwent at least two genome duplication events (Jiao et al. 2011), and subsequent multiple genome duplication events occurred in different groups of flowering plants. Genome duplication has major consequences that manifest both rapidly and over evolutionary timescales. Changes in gene expression patterns occur soon after genome duplication. For example, in newly formed allotetraploids of *Arabidopsis arenosa* and *A. thaliana* approximately 5% of genes were expressed at different levels than in the parental lines (Wang et al. 2006). In recently formed *Tragopogon* allopolyploids, approximately 76% of homoeologous genes displayed additive expression (the average of expression measured in each parental line), and approximately 20% of transcripts exhibited non-additive expression in which gene expression levels varied from the average of parents (Boatwright et al. 2018). Over longer time scales, a tendency of homoeologous gene expression from one parental genome to dominate over the other parental has been observed in several polyploids, including cotton, Brassicas and maize (Wendel et al. 2018). Re-diploidization by gene loss and recombination generally occurs after genome duplication, leading to differing contributions of the original genome complements to gene function (Woodhouse et al. 2014). Consequences of re-diploidization include the stable maintenance of functionally redundant copies, the rapid loss of genes, and gene neo-functionalization (Mandáková and Lysak 2018). Many crop species have undergone relatively recent polyploid events prior to and during domestication (Salman-Minkov et al. 2016). Understanding the genomic consequences of polyploidy in crops is centrally important, as hybrids are widely used to introduce new genetic variation into depleted crop plant gene pools and because they often exhibit superior vigour and yields compared to their parents. The molecular bases for this heterosis are currently not well understood (Birchler et al. 2010), with a variety of hypotheses proposed ranging from complementation of differing alleles, to misregulation of gene expression, to epigenetic explanations (Chen 2010; Ding and Chen 2018). For example, maize is an ancient tetraploid that has been re-diploidized by recombination of homoeologous chromosomes, leading to traces of two sub-genomes called maize1 and maize2, which differentially retain duplicated genes (Gaut and Doebley 1997; Schnable et al. 2011). These preferentially retained genes were more highly expressed (Renny-Byfield et al. 2017), and had different patterns of DNA methylation that were correlated with expression differences, but it was not possible to establish causal relationships between methylation differences and gene expression. Altered activities of transposable elements (TEs) and their patterns of DNA methylation have been proposed to underlie biased gene expression (Wendel et al. 2018). Analyses of TEs surrounding genes with biased gene expression were carried out in *Brassica rapa*, which has three sub-genomes that are re-diploidising (Cheng et al. 2016). These showed TE differences among the sub-genomes that were thought to be maintained from parental genomes, and there was a correlation between TEs and small RNAs located in putative promoter regions with dominantly expressed genes.

The wheat group of the Triticeae is characterised by stable tetraploid and hexaploid species that exhibit greater diversity, adaptability and potential for domestication than their diploid progenitors. Multiple types of genomic changes, including altered expression patterns of genes and TEs, and epigenetic changes, have been proposed to contribute to this “genomic plasticity” (Feldman and Levy 2015). The hexaploid bread wheat genome *(Triticum aestivum)* arose from integration of the diploid *Aegilops tauschii* DD genome into a tetraploid *T. turgidum* AABB genome less than 0.4 mya (million years ago) (Marcussen et al. 2014). The three component genomes are very closely related, sharing common ancestry in the Triticeae lineage approximately 6.5 mya. These three genomes behave as diploids at meiosis due to the pairing locus *Ph1* that limits pairing to homologous chromosomes during meiosis (Martín et al. 2017). The bread wheat genome is therefore a potentially informative experimental system for studying the interactions between closely related homoeologous loci in a stable genomic framework. Analyses of expression patterns of 1:1:1 syntenic homoeologous AABBDD “triads” in hexaploid wheat showed that 70% of triads exhibited balanced expression, in which each homoeolog was expressed at approximately the same level (Ramírez-González et al. 2018). The remaining triads displayed genome-biased and non-additive patterns of differential expression. In newly synthesised allohexaploid wheat (Chagué et al. 2010) and tetraploid AADD and S’S’A (where S’S’ is *Ae. longissima)* (Jiao et al. 2018) wheat lines between 60-80% of genes were additively expressed, suggesting a dynamic re-adjustment of gene expression patterns as a consequence of allopolyploidy. Rapid asymmetric changes in short RNA, histone methylation and gene expression in the two allotetraploids were also observed and were thought to contribute to genome-biased gene expression and the activation of TE transcription (Jiao et al. 2018). RNAseq analyses of newly formed allotriploid ABD and stable allohexaploid AABBDD lines generated from *T. turgidum* AABB with *Ae. tauschii* DD also showed rapid and extensive changes in gene expression in triploid tissues that were partly restored upon genome duplication (Hao et al. 2017).

Recent progress in assembling and annotating large and complex Triticeae genomes (Luo et al. 2017; International Wheat Genome Sequencing Consortium (IWGSC) et al. 2018) provides new opportunities for detailed spatial analyses of gene expression of Triticeae chromosomes in diploid and polyploid contexts. Our analyses of gene expression in homologous chromosome arms of *Ae. tauschii* and bread wheat show a chromosome-arm wide pattern of reduced expression in the hexaploid compared to the diploid genome context and multiple instances of new patterns of tissue-specific expression. Only a small proportion of these major differences could be related to structural changes in genes or altered patterns of gene methylation. In contrast, dramatically reduced chromatin accessibility in genes in the hexaploid context indicated a novel role for chromatin dynamics in altered gene expression.

## Results

### Assembly and annotation of wheat chromosome 3DL

We selected the long arm of chromosome 3 of *Ae. tauschii* and bread wheat for comparison as group 3 chromosomes of pooid grasses have not undergone chromosome fusion during their evolution (Luo et al. 2009; El Baidouri et al. 2017) and therefore have extensive conserved gene order and definable homoeologous relationships. A long-range assembly of *Ae. tauschii* AL8/78 has recently been constructed (Luo et al. 2017). We assembled chromosome 3DL from *T. aestivum* landrace Chinese Spring using a combination of sequenced Bacterial Artificial Chromosomes (BACs, Additional File 1) from a physical map (Lu et al. 2018), Triticum 3.1 assemblies of 3DL generated by PacBio sequencing (Zimin et al. 2017) and Fosill long mate pair reads (Lu et al. 2018). The order and orientation of the resulting 504 long scaffolds was assessed by alignment with the IWGSC v0.4 assembly of chromosome 3D (International Wheat Genome Sequencing Consortium (IWGSC) et al. 2018). The total assembly length of the chr3DL pseudomolecule was 371,759,578 bp (Table 1), similar to that of the IWGSC wheat assembly. The BAC and PacBio scaffolds corrected 19 orientation discrepancies in the IWGSC v0.4 Chinese Spring assembly. Although the IWGSC v1 assembly resolved 6 of these orientation discrepancies, it also introduced a long 25.37Mb orientation discrepancy.

**Table 1.** Sequence assembly of wheat chromosome 3DL

| Scaffolds | Versions | Number | Total bases<br>(n50) | n50 (bp) | Max (bp) |
| --- | --- | --- | --- | --- | --- |
| 3DL PacBio | Original | 2,703 | 409,078,553 | 279,970 | 1,657,081 |
|  | Fosill-joined | 1,751 | 422,542,707 | 437,268 | 2,142,379 |
|  | BAC-scaffoldable | 1,339 | 357,619,806 | 442,952 | 2,142,379 |
|  | BAC-joined | 524 | 371,055,817 | 1,096,174 | 5,859,701 |
| 3DL BACs | Original (v2) | 35,316 | 782,691,342 | 100,179 | 281,751 |
|  | PacBio-joined (v6) | 504 | 371,709,278 | 1,176,844 | 5,861,578 |
|  | pseudomolecule | 1 | 371,759,578 | - | - |

Fig. 1A shows alignments of the wheat 3DL, *Ae. tauschii* 3L and two versions of the IWGSC 3DL pseudomolecules (International Wheat Genome Sequencing Consortium (IWGSC) et al. 2018). There was essentially full-length alignment of the BAC-derived wheat 3DL pseudomolecule with this recently available wheat assembly and with *Ae. tauschii* chromosome arm 3L. Genes and pseudogenes were annotated with manual curation of transcript and predicted protein evidence. A total of 3,927 genes were identified of which 3,540 were located on anchored scaffolds. Seventy-five percent of the annotated genes were predicted with high confidence and 192 pseudogenes were identified based on gene models with conserved exon-intron structures that lacked an identifiable coding sequence. Repeats were identified on chromosome 3DL and 3L using RepeatMasker and annotated using repeat databases. Approximately 70% of the assembled sequences were classified as repetitive Additional File 2: Table S1), which comprised 52% class I LTR retrotransposons and 12% class II DNA transposons. Additional File 3 describes the assembly of chromosome 3DL and its relation to chromosome 3L of *Ae. tauschii (Luo et al. 2017).*

**Figure 1.**
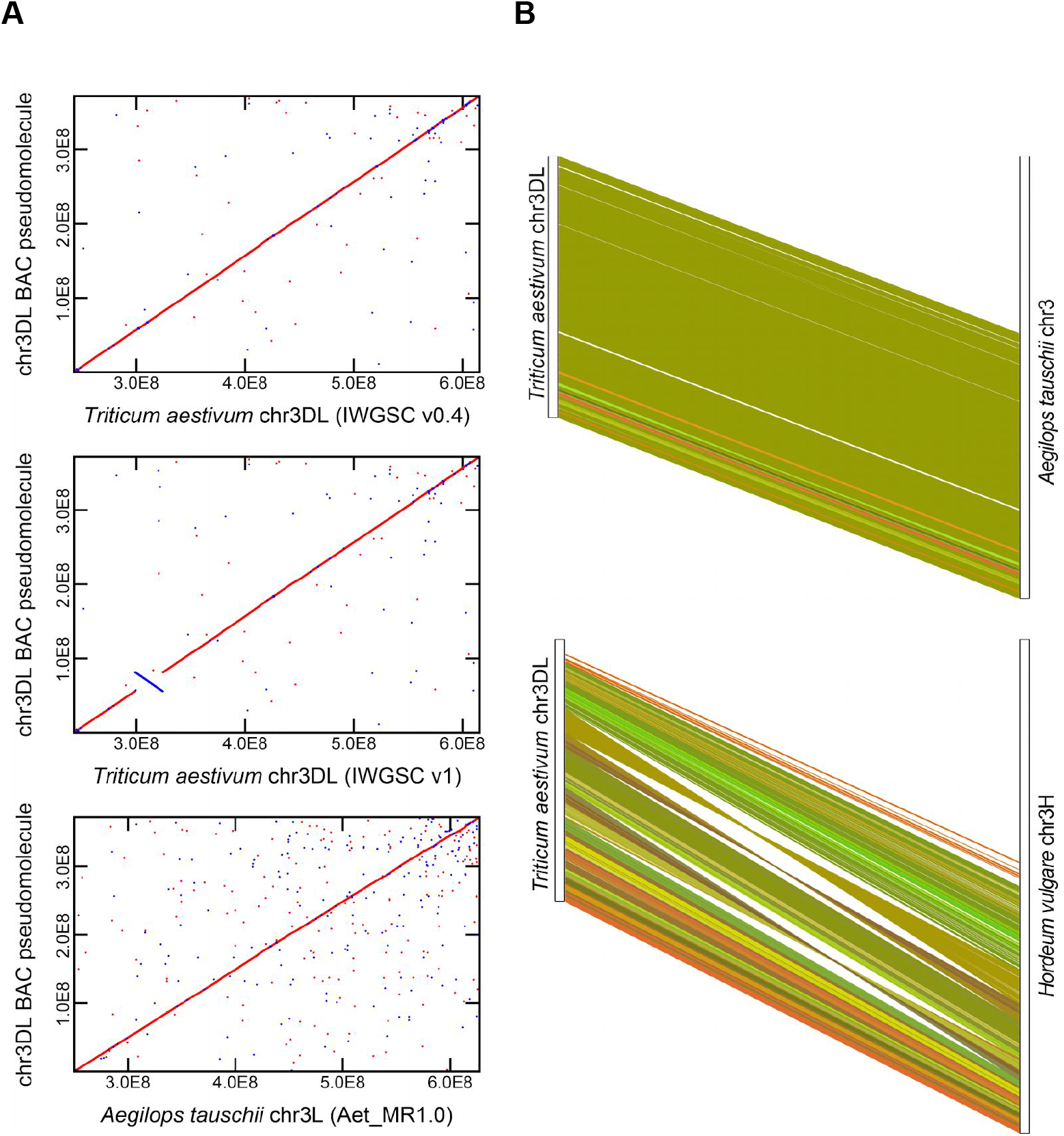
Chromosome alignments and gene synteny of the long arm of hexaploid wheat chromosome 3. **A**. The three panels show MUMmerplot alignments of the chromosome 3DL BAC-based pseudomolecule (on the y axis) to an early version (v0.4, top panel) and later version (v1) of IWGSC Chinese Spring chromosome 3DL and *Aegilops tauschii* AL8/78 chromosomes 3L on the X axes. Chromosome coordinates are in base-pairs, with chromosome 3DL BAC pseudomolecule coordinates starting at a centromeric location and extending to the telomere. Alignments are shown by the diagonal red line. The middle panel shows a large inversion (blue line) in the IWGSC v1 assembly of 3DL compared to the IWGSC v0.4 and BAC-based pseudomolecule. The lower panel shows an essentially collinear relationship between the sequence assemblies of wheat 3DL and *Ae. tauschii* homologous chromosome arms. **B**. Gene synteny alignments between 3,927 hexaploid wheat chromosome 3DL genes and 4,121 *Ae. tauschii* 3L genes. The similarities and collinearity of these genes and barley homologs was determined using BLAST+ and MCScanX. The coordinates of 3,456 gene pairs are shown by connecting lines between the chromosome arms. The different colours show different syntenic groups. The white region in the comparison of wheat and *Ae. tauschii* (upper panel) is due to an extended gene-free region. Contrast highly conserved gene order between wheat and *Ae. tauschii* with the loss of synteny between barley and hexaploid wheat.

### Conserved gene order between 3L and 3DL pseudomolecules

In order to make precise comparisons of *Ae. tauschii* and wheat, gene annotations were manually curated to similar standards on both chromosomes. The annotation of 3L (Luo et al. 2017) was extended by identifying a further 1,266 genes on 3L using *Ae. tauschii* AL8/78 RNAseq data and sequence similarity to 3DL gene models. This identified a total of 4,121 genes on 3L. Gene collinearity between wheat 3DL and *Ae. tauschii* 3L was defined by similarity searches and mapped using MCScanX (Additional File 4). Figure 1B shows extensive conserved syntenic gene order between the two chromosome arms, in which 3,456 of the 3,927 predicted 3DL wheat genes align to 3L genes with an average of 99.66% sequence identity across their full coding sequences (CDS; Additional File 4). Most of the 207 non-syntenic genes on wheat 3DL were located towards the telomeric end of the chromosome arms, suggesting that enhanced recombination at chromosome ends (See et al. 2006) may contribute to reduced synteny. The extensive collinearity of genes on 3DL with 3L contrasts with reduced gene order (due to three inversions) and gene similarity seen in comparison with barley (Fig. 1B). These alignments were used in subsequent comparisons of the two chromosome arms.

### Gene expression patterns in 3L and 3DL

RNAseq data was generated from RNA isolated from greenhouse grown *T. aestivum* Paragon and *Ae. tauschii* AL8/78 plants. RNA was isolated from leaves and roots of 2 week old plants, from 4 day old seedlings, and from 10 and 27 days after pollination (DAP) developing grain (Additional File 2: Table S2). Illumina TruSeq mRNA libraries were made from three independent polyA+ RNA preparations, and between 40M-90M reads were generated from each library. HiSAT2 (Kim et al. 2015) was used to map transcripts to each complete genome, and Stringtie (Pertea et al. 2016) was used to count reads and calculate transcripts per million (TPM) values (Additional File 4). Figure S1 in Additional File 2 shows the expression profiles of 3DL and 3L genes in the five sampled tissues. In total, 2,375 (68.72%) of the syntenic genes from 3DL (1,893 genes) and/or 3L (2,217 genes) were expressed at TPM≥1 in the five tissues examined (Additional File 4). Most genes were expressed in two or more tissues, while only ~5 % were specific to the tissues sampled. This pattern of substantial similarity of expression patterns from the two chromosome arms established a conserved framework to study quantitative differences in gene expression in the diploid and hexaploid contexts. Comparison of TPM values of syntenic gene pairs between hexaploid 3DL and diploid 3L (Fig. 2A) showed that approximately 70% of the pairs of genes showed a reduction of TPM values in the hexaploid 3DL context to approximately 40% of those measured in diploid 3L genomic context. This pattern of gene expression difference, which we call proportionately reduced expression, is shown as grey bars in Fig. 2B that mapped across the chromosome arms of wheat 3DL and *Ae. tauschii* 3L. TPM values are normalised to allow comparison of transcript levels within one genome. Quantitative comparison of gene expression in diploid and hexaploid gene complements is possible as both are normalised to the same factor that is independent of gene numbers. To validate the use of TPM for comparing gene expression between hexaploid wheat and diploid *Ae. tauschii*, we measured the expression of a set of 14 genes present as single copies in the AA, BB, DD and *Ae. tauschii* DD genomes (Additional File 5) using RNA extracted from 50,000 protoplasts and quantitative RT-PCR with a standard curve of a plasmid molecule. For 11 of the 14 gene sets, absolute transcript levels showed the same pattern of reduction in expression in the 3DL hexaploid context to approximately 40% of that measured in the D genome diploid context (Fig. 2C). The sum of A, B and D expression values was 1.2 times the diploid value on average (Fig. 2C). We therefore used TPM values for comparing gene expression values in diploid and hexaploid genome contexts.

**Figure 2.**
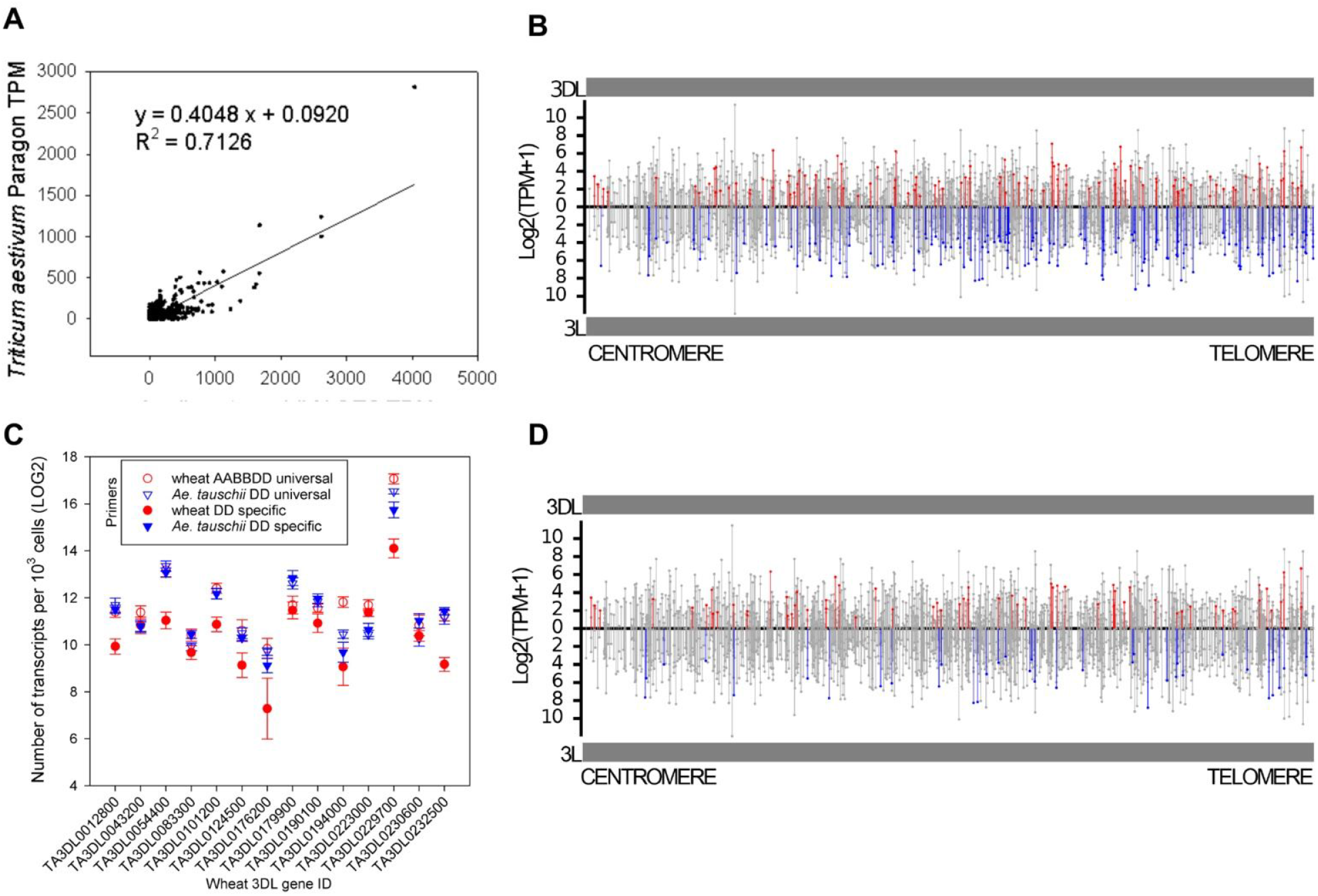
Comparison of expression of syntenic gene pairs in wheat 3DL and *Ae tauschii* 3L. **A** Comparison of gene expression levels between syntenic pairs of hexaploid Paragon 3DL and diploid AL8/78 3L genes. 2,378 (68%) of the syntenic genes were expressed in each of the five tissues examined. A trend of reduction of hexaploid wheat 3DL gene expression to 40% of that observed in diploid *Ae. tauschii* was observed using Transcripts Per Million (TPM) measures. **B** Chromosomal locations of differentially expressed and proportionately reduced genes in leaf tissue of Paragon 3DL and *Ae. tauschii* AL8/78. The locations of 3,456 syntenic gene pairs is shown on the horizontal axis, with the centromere on the left and telomere on the right. Gene expression values (TPM) are on the vertical axis, with wheat gene expression values shown on the upper panel and *Ae. tauschii* gene expression values shown on the lower panel. Differentially Expressed Genes (DEGs) were defined as those with ≽4-fold expression in wheat compared to *Ae. tauschii* (red lines), and ≽4 fold higher in *Ae. tauschii* compared to wheat (blue lines). Gene expression values of proportionately reduced genes (40% lower TPM in wheat compared to *Ae. tauschii)* are shown as gray lines. **C** Measuring expression levels of 14 syntenic gene pairs on wheat 3DL and *Ae. tauschii* 3L using absolute quantitative RT-PCR. The graph shows transcript levels expressed per leaf mesophyll protoplast on the vertical axis. The 14 genes (identified on the horizontal axis) were from among proportionately reduced genes and were selected based on the design of primer pairs for qRT-PCR that could distinguish all three AA, BB and DD homoeologs, and primer pairs that could specifically amplify only the DD gene copies in wheat and *Ae. tauschii.* For 11 of the 14 gene families, lower expression levels (41.78% on average) of the DD gene from hexaploid Paragon wheat were seen compared to the same gene in diploid *Ae. tauschii.* **D** Chromosomal locations of 106 conserved DEGs and balanced genes in leaf tissue of Paragon 3DL and three *Ae. tauschii* accessions AL8/78, Clae23 and ENT336. As described in panel **B** above, red lines mark those DEGs expressed more highly in wheat than the three *Ae. tauschii* varieties, and blue lines indicate DEGs with higher expression in the three *Ae. tauschii* lines compared to Paragon wheat. Gray lines indicate proportionately reduced gene expression patterns.

### Differentially expressed genes

As described above, approximately 70% of the 3DL-3L gene pairs showed proportionately reduced expression in wheat compared to *Ae. tauschii.* Another general pattern of changes in gene expression was also observed; many genes showed large differences in expression, with some genes expressed much more in *Ae. tauschii* than wheat, and *vice versa.* To define a set of these differentially expressed genes (DEGs), we first normalized wheat TPM values to *Ae. tauschii* TPM values by multiplying them by a factor (1.7604) to take account of the 70% of genes that had reduced expression levels in 3DL compared to 3L (Fig. 2A). After this normalisation, a threshold value of ≥ 4-fold change (either up in wheat or up in *Ae. tauschii)* was established as a conservative measure, for example compared to 30% variation used in whole genome analyses (Ramírez-González et al. 2018). This defined a class of 674 DEGs (28.38% out of the 2,375 expressed genes) in the 5 examined tissues. Their numbers are shown (Additional File 2: Table S3) and represented as red bars for those with ≥4-fold increase in wheat or blue bars for ≥4-fold increase in *Ae. tauschii* on a chromosome map (Fig. 2B). It is possible that these differences in expression between wheat and *Ae. tauschii* were due to differences between the sequenced *Ae. tauschii* variety AL8/78 and the currently unknown donor of the D genome to bread wheat. This is a valid consideration as AL8/78 cannot form a stable hybrid with tetraploid wheat lines (unpublished observations). We therefore carried out RNAseq analyses of two other diverse *Ae. tauschii* varieties, Clae23 and ENT336, which can form viable hybrids with tetraploid wheat (unpublished observations). Leaf RNAseq data from Clae23 and ENT336 was mapped to 3L (Additional File 2: Figure S2a, b). Of the 262 (16% of 1,564 expressed genes) leaf-specific DEGS identified in wheat 3DL and *Ae. tauschii* AL8/8 3L, 106 were conserved in all three *Ae. tauschii* varieties. Mapping of these conserved DEGs to 3DL and 3L showed that they occurred along the entire chromosome arm, with no evidence for regional differences (Fig. 2D).

Gene sequences (exon and UTR) were highly conserved between wheat 3DL and *Ae. tauschii* 3L, with some reduction in similarities towards their telomeres (Fig. 3A). Similarly, promoter sequence variation was also more pronounced towards the telomeres, but no clear relationships between promoter and gene sequence divergence and differential patterns of gene expression were seen (Fig. 3B). For example, only 4 of the 106 leaf DEGs showed minor differences in exon-intron structure (Additional File 2: Figure S3).

**Figure 3.**
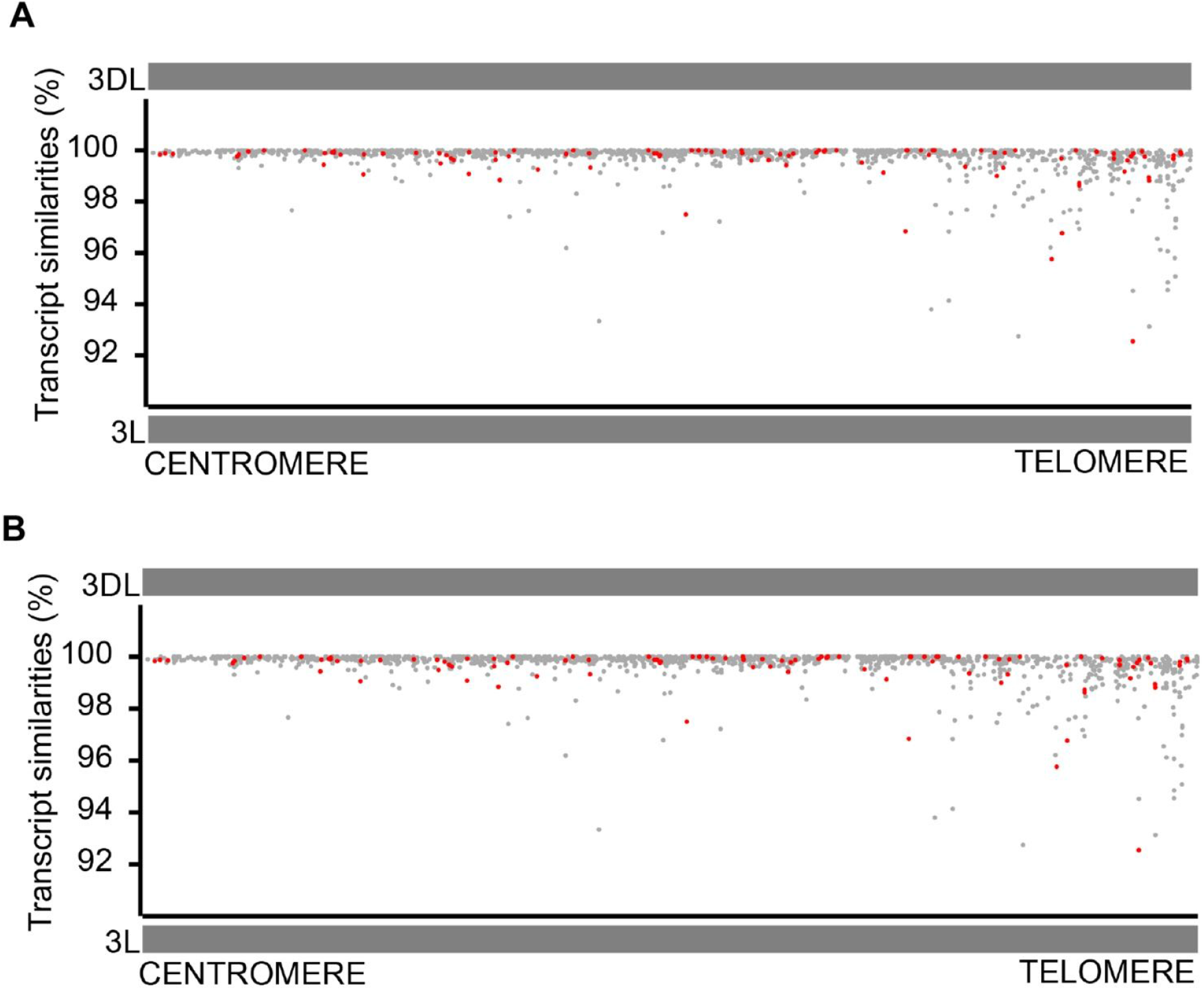
Sequence similarities of transcripts and promoters of 106 wheat 3DL syntenic gene pairs expressed in three *Ae. tauschii* lines. **A** Sequence differences between gene transcripts of 106 syntenic genes with conserved differential gene expression patterns on the long arms of wheat chromosome 3DL and 3L of *Ae. tauschii.* The vertical axis shows % similarities in predicted transcript sequences. Red dots indicate the 106 consensus DEGs among 3 *Ae. tauschii* accessions, and the gray dots identify genes with proportionately reduced expression. Increased transcript sequence divergence is seen towards the telomere end of the the chromosome arm, while DEGs and gene with proportionately reduced expression were distributed along the chromosome arm. **B** Sequence differences between promoters of 106 syntenic genes with conserved differential gene expression patterns on the long arms of wheat chromosome 3DL and 3L of *Ae. tauschii.* The vertical axis shows % similarities in predicted promoter sequences, defined as 2kb upstream of the predicted transcription start site. Red dots indicate the 106 consensus DEGs among 3 *Ae. tauschii* accessions, and the gray dots identify genes with proportionately reduced expression. Promoter sequence divergence was seen along the chromosome arm, but there was no clear relationship between promoter divergence and differential gene expression.

DEGs were most frequently identified in RNA isolated from 27 DAP developing grain of wheat and *Ae. tauschii.* Among the 86 up-regulated 3DL genes in 27 DAP developing grain of Paragon hexaploid wheat, 63 were functionally classified using GO terms as involved in protein targeting and degradation, RNA transcription, processing and translation regulation, and several motifs were enriched in the promoter regions of these genes, suggesting the DD genome contributes genes with new roles in the late-stage of developing grain (Additional File 6). When these 86 DEGs were mapped to TGAC v1 chr3DL genes (Clavijo et al. 2017), 33 had accompanying tissue-specific gene expression data. Twenty-seven of these 33 genes were mostly highly expressed in 20 DAP aleurone layer samples. The 33 genes were also expressed in 20 DAP whole endosperm, 20 DAP transfer cell, and 20 and 27 DAP starchy endosperm tissue samples. These analyses show that upon formation of the hexaploid genome, 3L genes are subject to differential regulation during later stages of grain development.

### Comparative DNA Methylation Analyses

Differences in DNA methylation, perhaps mediated by repeat elements, have been proposed to confer differences in gene expression between the subgenomes of polyploids (Wendel et al. 2018; Wang et al. 2004). To assess the potential involvement of gene body and promoter methylation in the two general patterns of gene expression differences observed in chromosomes 3L and 3DL, we conducted exome capture and bisulphite sequencing of *Ae. tauschii* AL8/78, and whole genome bisulphite sequencing of Paragon hexaploid wheat DNA. Reads were mapped to the complete genomes, those mapping to chromosomes 3L and 3DL were defined, and methylation in CpG, CHG or CHH contexts was identified. Overall levels of 3L and 3DL methylation were similar in wheat 3DL and *Ae. tauschii* 3L chromosome arms, with 89.9%, 59.4% and 3.8% methylation of CpG, CHG and CHH sites for wheat and 87.1%, 53.4% and 3.4% for *Ae. tauschii* AL8/78 (Additional File 2: Table S4). To compare average methylation levels across promoters and gene bodies for 3DL and 3L genes, their promoter and gene lengths were normalised. First, average methylation levels were assessed across the promoters and gene bodies of expressed and non-expressed genes for CpG, CHG and CHH contexts to define relationships between gene methylation and gene expression. Fig. 4 shows a decrease of CpG methylation at the Transcriptional Start Site (TSS) and Transcription Termination Site (TTS) in comparison to the promoter and gene body regions. This pattern of reduced TSS methylation of CpG and CHH contexts is more marked for expressed genes than non-expressed genes in both 3L and 3DL genes. CpG and CHG methylation at the TSS (+/− 20bp) is significantly lower in expressed genes compared to non-expressed genes (CpG sites; p < 0.0001, t=5.9739, df=80, CHG sites; p< 0.0001, t=6.3446, df=80).

**Figure 4.**
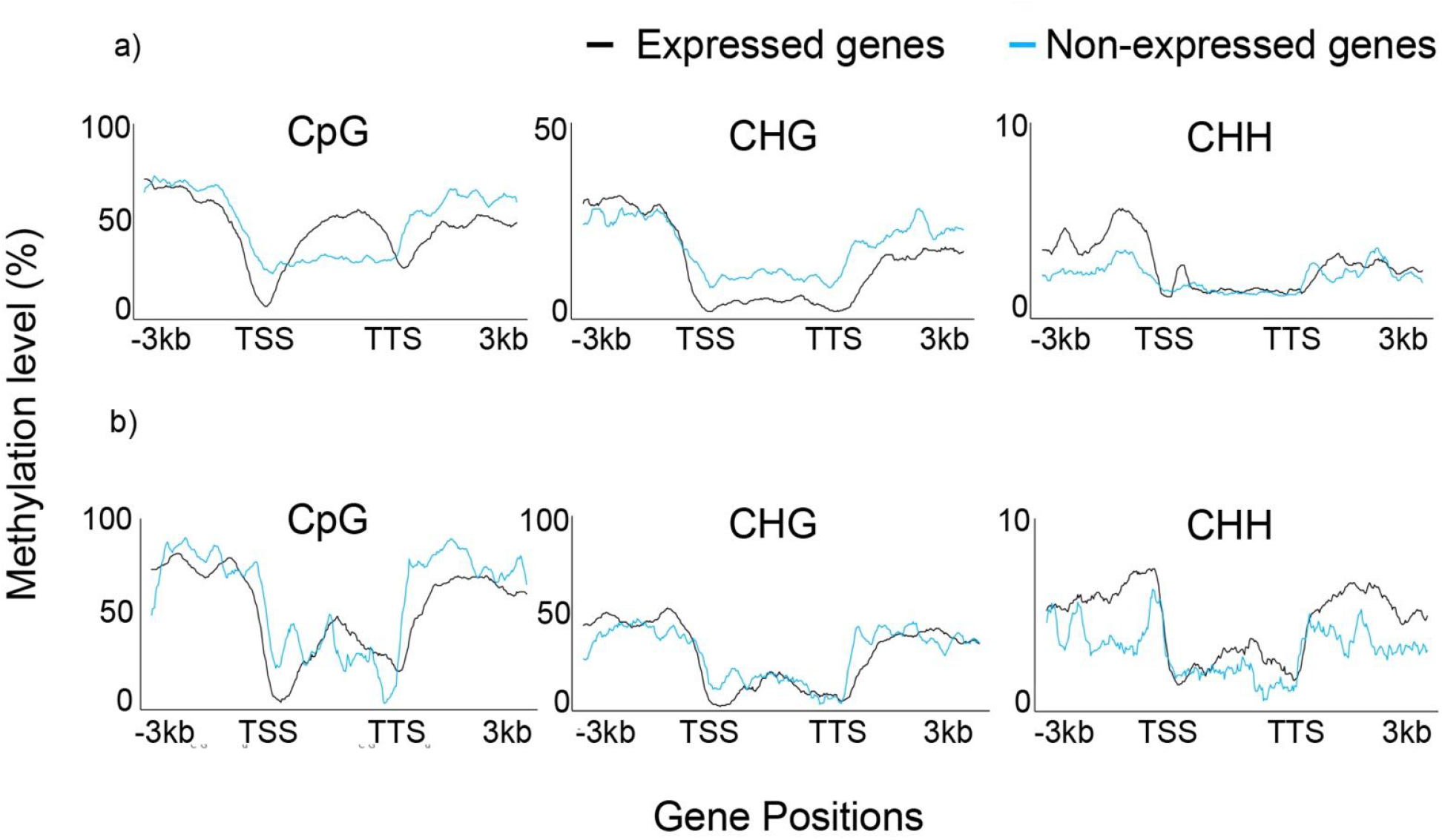
Average methylation across all expressed and non-expressed genes on hexaploid wheat 3DL and diploid *Ae tauschii* 3L chromosome arms. **A** The distribution of CpG, CHG and CHH DNA methylation contexts across expressed and non-expressed genes on wheat chromosome 3DL. TSS: Transcriptional Start Site; TTS: Transcriptional Termination Site. Methylation was assessed 3kb upstream of TSS and 3kb downstream of the TSS. **B** The distribution of CpG, CHG and CHH DNA methylation contexts across expressed and non-expressed genes on *Ae. tauschii* AL8/78 chromosome 3L. TSS: Transcriptional Start Site; TTS: Transcriptional Termination Site. Methylation was assessed 3kb upstream of TSS and 3kb downstream of the TSS.

Second, we compared gene-body and promoter methylation patterns between 3DL and 3L genes. For this analysis, DNA methylation for each of three contexts (CpG/CHG/CHH) was averaged independently across each gene-body and promoter region. A gene/promoter region was only analysed if a minimum of 5 methylated cytosines were included in the region, each with a minimum bisulphite sequencing coverage of 5X for *Ae. tauschii* 3L and lOX for Paragon 3DL, as it had a higher sequence depth coverage. This yielded a total of 2,709 unique gene-regions (81.9% of the total genes) and 2,182 unique promoter regions across the CpG/CHG/CHH contexts for chromosome 3DL and 2,952 unique gene-regions (71.5% of the total genes) and 2,719 unique promoter regions for chromosome 3L (Additional File 2: Table S5). Alignments between 3,456 gene pairs on 3DL and 3L permitted comparison of methylation between Paragon and *Ae. tauschii* for 2,224 gene pairs in total (64.35%) across CpG/CHG/CHH gene and promoter sites (1,920 genes and 1,368 promoters). From this comparison differentially methylated genes (DMGs) were identified. A DMR was called if a CpG region showed a difference in methylation of 50% or more (q value < 0.05), a CHG region showed a difference of 25% or more or a CHH site showed a difference of 10% or more. Overall, only 11.3 % of differentially methylated genes or promoters were associated with differences in gene expression between Paragon 3DL and *Ae. tauschii* 3L (Table 2). This suggested that differential promoter and gene methylation potentially accounts for a small proportion of observed gene expression differences between diploid *Ae. tauschii* 3L and hexaploid wheat 3DL genes.

**Table 2.** Number of differentially methylated genes that are also differentially expressed between wheat 3DL genes and *Ae tauschii* 3L genes

|  | **CpG** | **CHG** | **CHH** |
| --- | --- | --- | --- |
| Genes showing differential methylation (q< 0.05) | 526 | 169 | 42 |
| Genes showing differential methylation & DEGs | 64 | 18 | 5 |
| Promoters showing differential methylation (q< 0.05) | 263 | 85 | 123 |
| Promoters showing differential methylation & expression 4-fold | 27 | 6 | 17 |

### Pseudogene Analyses

Gene loss as a consequence of polyploidisation has been proposed to be an important driver of genomic change, and pseudogene formation is an important contributor (Wicker et al. 2011). A total of 192 genes on chromosome 3DL were classified as pseudogenes, based on well defined exon-intron structures using EST, protein and RNAseq evidence, with predicted introns obeying the GT-AG rule, but without a predictable coding sequence (CDS) region. 176 of these predicted pseudogenes were anchored on the chr3DL pseudomolecule and 16 were on unlocalised scaffolds. Among the 160 pseudogenes that were syntenic on 3DL and 3L, 66 (43 in Paragon and 49 in AL8/78) were expressed with at least 1 TPM in the examined tissues. DNA methylation analyses of the 160 syntenic pseudogenes revealed CpG, CHG or CHH gene methylation data for 85, 82 and 101 genes and 45, 46 and 67 promoters respectively. Table 3 compares the methylation levels of pseudogenes with the methylation patterns of all analysed genes on 3L and 3DL. In general methylation levels were elevated in pseudogenes in both Paragon and *Ae. tauschii*, by 13.2% for CpG sites and 12.6% for CHG sites, in comparison to non-pseudogenes. Methylation levels were not observed to increase at CHH sites in pseudogenes (average difference −0.4%). We identified seven pseudogenes on wheat 3DL that were intact genes in *Ae. tauschii* 3L, indicating a potential recent origin (Table 4). Three of these *Ae. tauschii* 3L genes were expressed, but none were in Paragon wheat, confirming their identification as pseudogenes. Available methylation data for 4 of these gene pairs showed a large increase in CpG methylation in the wheat pseudogenes compared to their functional *Ae. tauschii* counterparts. Only small changes in CHG and CHH methylation (Table 4) were identified in the wheat-specific pseudogenes.

**Table 3.** Methylation levels of pseudogenes in Paragon wheat 3DL and *Ae tauschii* AL8/78 3L

| **Methylation context** | **CpG %** | **CHG %** | **CHH %** |
| --- | --- | --- | --- |
| Paragon wheat 3DL |  |  |  |
| Coding genes | 59.7 | 9.3 | 1.2 |
| Coding gene promoters | 65.4 | 30.7 | 3.7 |
| pseudogenes | 72.5 | 25.2 | 1.4 |
| pseudogene promoters | 78.3 | 43.6 | 2.7 |
| *Ae tauschii* AL8/78 3L |  |  |  |
| Coding genes | 29.9 | 11.7 | 2.1 |
| Coding gene promoters | 62.5 | 38.1 | 5.5 |
| pseudogenes | 53.7 | 28.8 | 2.7 |
| pseudogene promoters | 65.7 | 42.7 | 4.0 |

**Table 4.** Pseudogenes in Paragon wheat 3DL that have intactl counterparts in *Ae. tauschii* AL8/78 3L

| Paragon GeneID | Gene Expression | Mutations | Descriptions |  | CpG % | CHG % | CHH % |
| --- | --- | --- | --- | --- | --- | --- | --- |
| TA3DL0067600 | - | Exon3 non-sense mutation (TAA/GAA) | MICOS complex subunit MIC60-like | - | - | - | - |
| TA3DL0149600 | 10DAP developing grain and seedling in AL8/78 | Exon2 Opal mutation (TGA/CGA) | Protein ACCELERATED CELL DEATH 6-like | Paragon | 76.97 | 0.88 | 0.17 |
|  |  |  |  | AL8/78 | 4.00 | 3.83 | 0.69 |
| TA3DL0184600 | 10 DAP and 27 DAP developing grain, root and seedling in AL8/78 | Exon4 2 bases insertion (GTGAG/GAG) | GDP-L-galactose phosphorylase 1-like | Paragon | 47.30 | 0.37 | 0.39 |
|  |  |  |  | AL8/78 | 45.71 | 8.89 | 1.48 |
| TA3DL0211300 | Root in AL8/78 | Exon4 non-sense mutation (TGA/GGA) | Endo-1,3;1,4-beta-D-glucanase-like | Paragon | 49.28 | 1.49 | 0.26 |
|  |  |  |  | AL8/78 | 12.00 | 3.64 | 1.39 |
| TA3DL0286700 | - | Exon5 one deletion (-TC/TTC) | Serine/threonine-protein phosphatase 7 long form homolog | Paragon | 97.54 | 66.33 | 1.49 |
|  |  |  |  | AL8/78 | 97.35 | 50.81 | 2.32 |
| TA3DL0299800 | - | Exon2 alternative splicing | Uncharacterized | - | - | - | - |
| TA3DL0320700 | - | Exon4 Opal mutation (TGA/CGA) | Zinc finger MYM-type protein 1-like | - | - | - | - |

### Chromatin accessibility

Analyses of DNA methylation data identified gene body and promoter methylation differences in only 11.3% of genes that were differentially expressed between *Ae. tauschii* 3L and wheat 3DL (Table 2). This suggested that DNA methylation may make a relatively minor contribution to differential gene expression. What may then account for these major differences in gene expression patterns between the highly conserved diploid and hexaploid chromosome arms? The chromosome-wide patterns of differences in gene expression between 3L and 3DL are reminiscent of patterns of change in gene expression seen in dosage compensation (Giorgetti et al. 2016) and in large-scale transcriptional reprogramming (Miyamoto et al. 2018). In these examples differences in chromatin accessibility were key factors in altered gene expression patterns. We carried out ATAC-seq (Assay for Transposase-Accessible Chromatin) (Buenrostro et al. 2013) in nuclei prepared from leaf protoplasts of hexaploid Paragon and diploid *Ae. tauschii* AL8/78. 150bp paired-end reads from three independent ATAC assays were mapped to each complete genome and those mapping to 3DL and 3L were analysed. Four classes of DNA sequence were assessed for chromatin accessibility: 5’UTR + promoter (including from the ATG to 2kb upstream), CDS (the gene coding region and predicted intron sequences), 3’UTR + downstream (including the 3’ UTR to 2kb downstream), and intergenic (regions >2kb distance from genes). ATAC peak areas in these classes of DNA sequence were identified by MACS2 (Zhang et al. 2008), and only peak areas present in all three independent experiments were used in subsequent analyses. Sequence reads mapping to the chloroplast genomes of wheat and *Ae. tauschii*, and the mitochondrial genome of wheat, were identified and removed as these are due to Nextera reactions with the DNA of these organelles, which contaminate nuclear preparations. This removed large spurious ATAC peaks associated with numerous chloroplast and mitochondrial insertions in each chromosome arm. Figure S4 in Additional File 2 shows the profile of ATAC fragment sizes on chromosomes 3L and 3DL. The ~10.5 bp periodicity reflects cleavage of the DNA helix, and a trace of single and double nucleosome spacing can be seen in the 3L ATAC fragment length frequency plot (Buenrostro et al. 2013). In both *Ae. tauschii* and Paragon wheat, normalised density plots of ATAC peaks showed that these were mainly found in 5’UTR + promoter regions, consistent with the demonstration of more accessible chromatin in putative transcriptional regulatory regions in mammalian cells (Buenrostro et al. 2013). Intergenic regions had the lowest ATAC peak densities, although the absolute numbers of peaks in intergenic regions were high (Additional File 2: Table S6 and Figure S5). Peak lengths showed that regions of accessible chromatin extended to over 5 nucleosomes, with a peak at 1-2 nucleosome spacing. In *Ae. tauschii* leaf nuclei 4,960 ATAC peaks on 936 genes were identified on chromosome 3L, compared to 2,970 ATAC peaks in 1,187 genes on chromosome 3DL of hexaploid wheat leaf nuclei (Additional File 2: Table S6). Interestingly, 26.53% of Paragon and 28.64% of *Ae. tauschii* ATAC peaks mapped to intergenic space. Approximately ⅔ of the intergenic ATAC peaks were in regions that were not annotated as repeats, while nearly ¾ of the ATAC peaks that mapped to annotated repeats were found over simple sequence repeats (SSR) (Fig. 5A). Nevertheless, only a small proportion of the annotated repeats on 3DL and 3L had accessible chromatin, the largest of which was 8% of SSR on 3L (Fig. 5B). The remaining ATAC peaks mainly covered both Class I LTR retrotransposons and Class II DNA transposons, with DNA repeats appearing to have more accessible chromatin. These data show that most repeat classes, in particular SSRs, have a small proportion of accessible chromatin.

**Figure 5.**
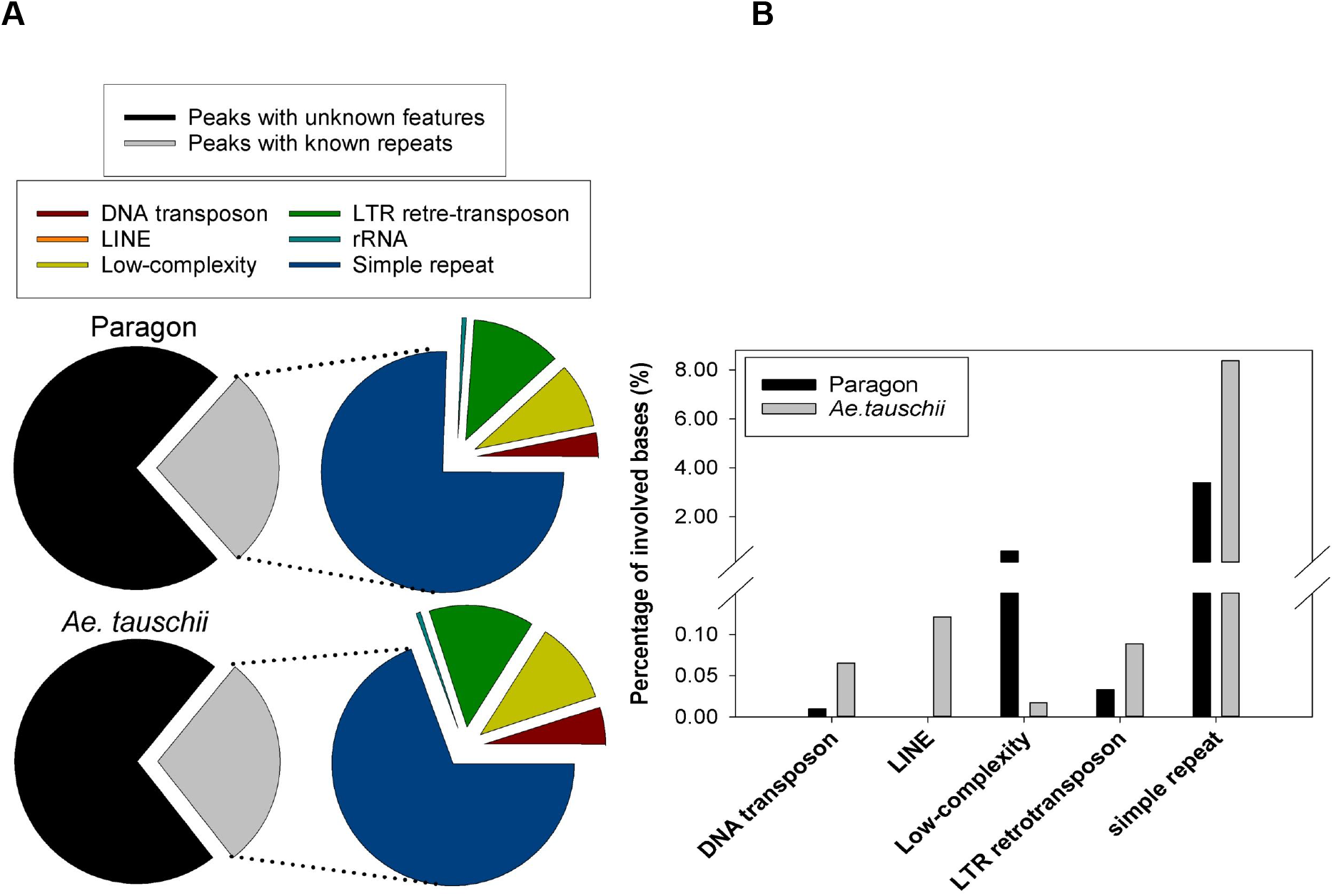
Distribution of chromatin accessibility across intergenic regions of hexaploid wheat 3DL and *Ae. tauschii* 3L chromosome arms. **A** The proportion of open chromatin sequences in intergenic regions. The left pie charts show the proportion of accessible chromatin sequences in intergenic space that has no annotated features and annotated repeats on both chromosome arms. The right pie charts show the proportion of different annotated repeats that have accessible chromatin. **B** The graph shows the percent of annotated repeats that have accessible chromatin in hexaploid wheat 3DL and *Ae. tauschii* 3L chromosome arms.

In contrast to these relatively similar patterns of chromatin accessibility in intergenic regions of *Ae. tauschii* 3L and wheat 3DL, major differences in patterns of chromatin accessibility were seen in the genic regions of *Ae. tauschii* 3L and Paragon 3DL. Peak length density plots (Fig. 6A) showed an overall restriction of chromatin peak lengths in genic regions of Paragon wheat 3DL compared to *Ae. tauschii* 3L. These differences were highly significant for genes with both proportionately reduced and differential patterns of of gene expression differences (Fig. 6B). Figs. 6C, 6D shows typical patterns of reduced ATAC peaks on the promoters of two syntenic pairs of wheat genes compared to their *Ae. tauschii* counterparts that involved loss of additional peaks and reduced width of a common peak. When peak length distributions were mapped to the Transcriptional Start Sites (TSS) of wheat 3DL and *Ae. tauschii* 3L genes, a clear pattern of more extensive chromatin accessibility was seen in 3L genes, with approximately 60% of accessible regions found within 100kb of the TSS. In contrast, 50% of accessible chromatin was found within 3kb of wheat 3DL gene TSSs (Fig. 6E). This large-scale restriction of genic chromatin accessibility in hexaploid wheat 3DL compared to *Ae. tauschii* 3L genes mapped across the entire chromosome arms (Fig. 6F; Additional File 2: Table S7).

**Figure 6.**
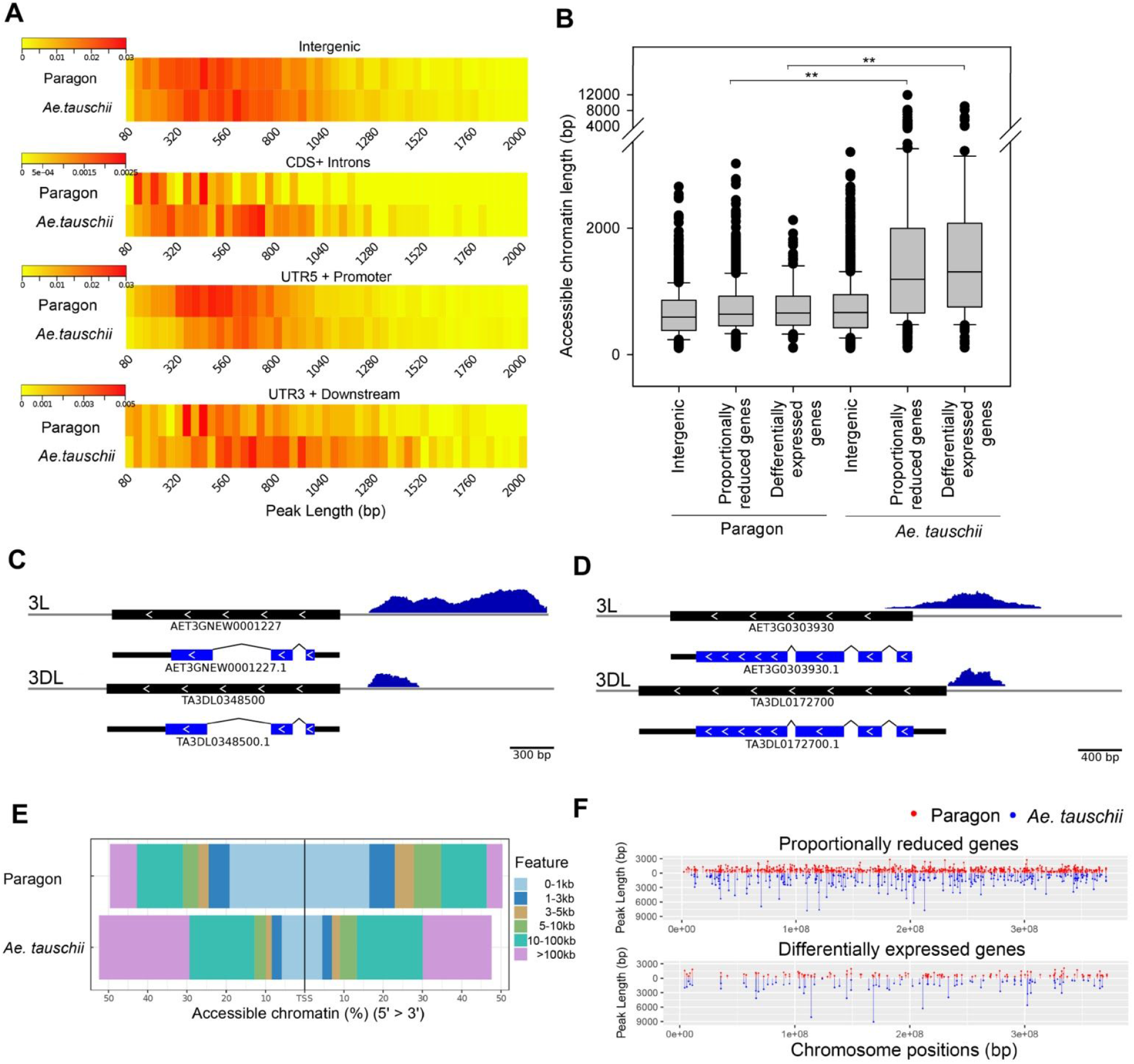
The distribution of accessible chromatin on wheat chromosome 3DL and *Ae. tauschii* 3L. **A** Normalized ATAC peak length enrichment for four classes of chromosomal regions on wheat chromosome 3DL and Ae. tauschii 3L. The chromosomal regions are intergenic; CDS+introns; 5’UTR+2kb upstream putative promoter region; 3’UTR+2kb downstream. ATAC peak length distributions are shown in base-pairs on the horizontal axis. The colour scale shows ATAC peak frequency distribution per bin. **B** Box plots of ATAC peak lengths across intergenic regions, genes with proportionately reduced expression patterns, and genes showing differential expression between wheat 3DL and *Ae. tauschii* 3L. The significance of ATAC peak length differences was assessed by one way ANOVA. Peak length differences for balanced genes between wheat at *Ae. tauschii* were significant at 6.99E-68, and 1.34E-10 for differentially expressed genes. **C** An example of ATAC peaks on a pair of *Ae. tauschii* and wheat genes with typical proportionately reduced gene expression. **D** An example of ATAC peaks on a typical pair of differentially expressed *Ae. tauschii* and wheat genes. **E** Normalised distance distribution of ATAC-sequence peaks relative to the TSS of genes in hexaploid wheat and diploid *Ae. tauschii* genes. **F** ATAC peak length distributions across syntenic genes with either proportionately reduced expression patterns (upper panel) or differential expression patterns (lower panel) across chromosome 3DL. Red lines and dots mark wheat ATAC peak lengths on the promoter+5’UTR regions of genes, while blue lines and dots mark ATAC peak lengths on the promoter+5’UTR regions of *Ae. tauschii* genes.

Of the 683 genes with differential ATAC peaks between wheat and *Ae. tauschii*, 133 of 159 genes (84%) that were differentially expressed in leaf tissue (the tissue used for ATAC-seq) also had differential ATAC peaks (Fig. 7). Of these, most different ATAC peaks occurred in the 5’UTR+promoter region. These analyses show that the majority of genes exhibiting proportionately reduced expression in wheat 3DL had reduced chromatin accessibility, and that DEGs also had varying ATAC peak distributions, mainly over their 5’ UTR5+promoter regions. These pervasive patterns of altered chromatin accessibility and gene expression indicate an important role for chromatin accessibility in gene expression in allopolyploid wheat.

**Figure 7.**
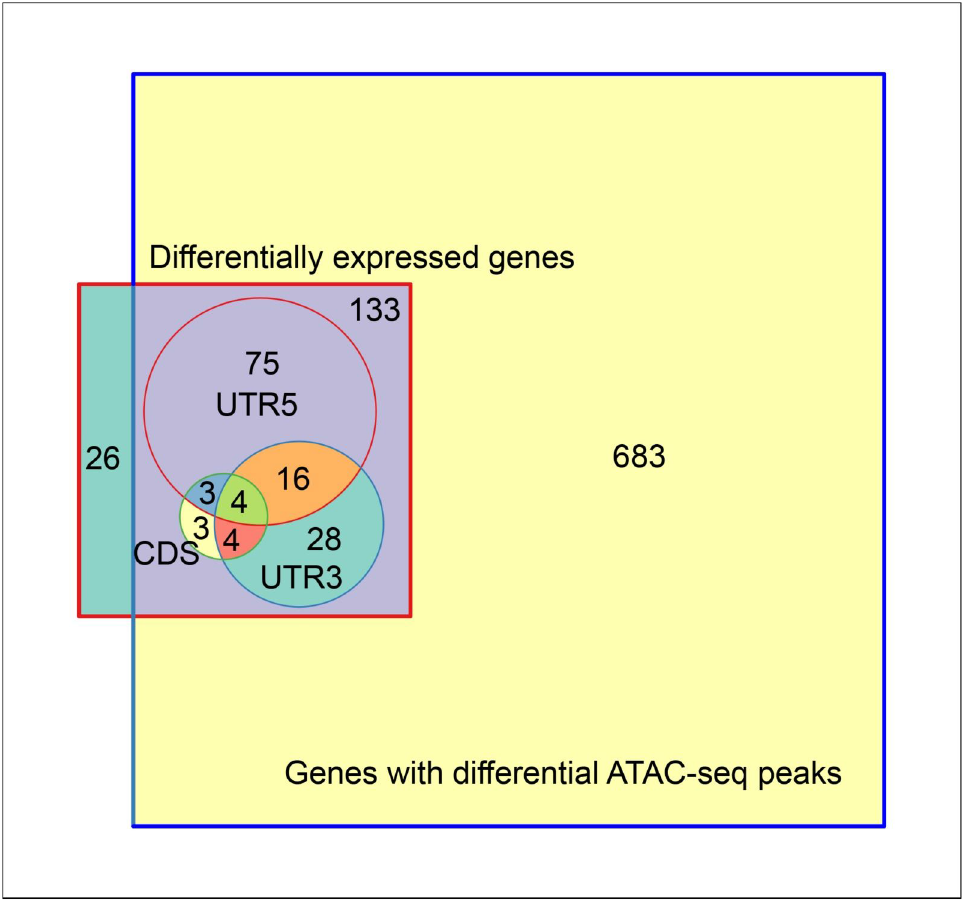
Relationships between differentially expressed genes and differential ATAC peaks. The Venn diagram shows 683 genes with differential ATAC peaks on any region of the gene between wheat and *Ae. tauschii.* 133 of 159 genes that are differentially expressed in leaf tissue (the tissue used for ATAC-seq) have differential ATAC peaks. Of these, most differential ATAC peaks occurred in the 5’UTR+promoter region (shown as UTR5).

## Discussion

Many studies have identified large-scale patterns of change in gene expression in polyploid plants compared to their progenitors (Chen and Ni 2006). We used long-range assemblies of homologous chromosome arms of diploid *Ae. tauschii* (3L) and hexaploid bread wheat (3DL) to characterise expression differences between essentially complete sets of syntenic genes from these chromosome arms in diploid and hexaploid genome contexts. The high degree of sequence conservation of the 3DL chromosome arm in hexaploid wheat and its progenitor diploid species *Ae. tauschii*, coupled to detailed annotation of genes and repeats, provided an extensive framework of nearly 3,500 gene pairs with 99.66% sequence similarity in their coding regions. Despite this extensive conservation of gene content on 3L and 3DL, gene expression patterns were radically different in the diploid and hexaploid genomic contexts. Nearly 70% of the gene pairs were expressed in the five analysed tissues (Additional File 2: Figure S1), and of these, 70% showed a proportionate reduction in gene expression in wheat, in which expression of the DD gene complement in the AABBDD genome fell to 40% of that measured in the DD genome of *Ae. tauschii*, and the overall expression of 1:1:1 AABBDD homoeologs was 1.2 times higher than in the DD genome. This pattern of reduced expression may reflect balancing gene expression in which approximately 70% of 1:1:1 AABBDD homoeologs are expressed at an approximate mid-point level (Ramírez-González et al. 2018). Similar overall reductions in expression have also been observed in newly formed wheat hybrids (Akhunova et al. 2010; Jiao et al. 2018; Chelaifa et al. 2013), suggesting a dynamic re-alignment of gene expression to near-diploid levels is an early consequence of allopolyploidy for the majority of wheat genes. The remaining 30% of gene pairs on 3L and 3DL exhibited differential expression levels that were significantly higher or lower in the hexaploid 3DL context compared to the diploid 3L context. This proportion is again similar to that observed in the complete wheat genome (Ramírez-González et al. 2018). We could not identify a clear general relationship between promoter sequence differences in 3L and 3DL gene pairs and differential gene expression patterns, and differences in exon-intron structures that may account for differential expression were discounted due to the very high similarity of gene pairs on 3L and 3DL. A significant proportion of DEGs were expressed during the later stages of grain development (Additional File 6). Notably, among the 86 DEGs that were up-regulated in developing hexaploid wheat grains compared to *Ae. tauschii*, many were involved in protein and RNA regulation, and of the 33 genes with a known tissue-specific expression patterns (Pfeifer et al. 2014), 27 were most highly expressed in the aleurone layer. This co-opting of 3L genes to wheat grain development through altered expression levels is consistent with the differential contributions of the AA, BB and DD genomes to functional modules involved in seed development identified by co-expression analyses (Pfeifer et al. 2014). Enrichment of promoter motifs in these 87 DEGs (Additional File 6) is consistent with a model in which transcriptional regulation from the AABB genomes may differentially regulate DD genes during later stages of grain development. Such emergent patterns of *cis-trans* interactions in polyploid genomes have been modelled (Bottani et al. 2018; Hu and Wendel 2018).

Differences in gene methylation have been proposed to contribute to differences in gene expression between progenitor and allopolyploid species (Chen and Ni 2006; Hu et al. 2013). To assess the role of gene methylation on expression patterns of 3L and 3DL genes, we carried out bisulphite sequencing of the complete hexaploid Paragon genome and of the gene space of *Ae. tauschii 3L.* Similar overall levels and patterns of CpG, CHG and CHH methylation were seen on both 3L and 3DL genes, and there was a general negative correlation between the extents of CpG and CHH methylation on TSS and gene expression (Table 2). Only 11% of DEGs had different gene and promoter methylation patterns that might cause altered expression, indicating a relatively small influence of methylation differences on gene expression patterns in 3DL and 3L. In contrast, of the seven pseudogenes found in 3DL that had an intact homolog in 3L, available bisulphite sequence data for four showed large increases in CpG methylation. This is consistent with extensive methylation of pseudogenes seen in many plant species (Schöb and Grossniklaus 2006; Meunier et al. 2005). Whether DNA methylation is a cause or consequence of pseudogene formation is not known in these examples.

Given the lack of evidence for differences in sequence composition (including gene loss and pseudogenization) or DNA methylation that could account for the major patterns of altered gene expression observed across chromosomes 3L and 3DL, what mechanisms may be responsible? Changes in small RNA and chromatin, as measured by chromosome immunofluorescence (Jiao et al. 2018), accompany the formation of new allotetraploid wheat lines, suggesting some forms of chromatin modification may contribute to altered gene expression in wheat allopolyploids. Such changes occur rapidly after polyploid stabilisation and are stable in newly formed wheat (Chagué et al. 2010) and Arabidopsis (Shi et al. 2012) allopolyploids. These observations suggest chromatin-based mechanisms could impose rapid, persistent and reversible genome-level changes on gene expression in allopolyploids in the absence of extensive changes in DNA methylation and gene loss.

Dynamic interplay between nucleosomes, transcription factors and chromatin remodelling proteins alters physical access to DNA in chromatin (Klemm et al. 2019). Physical access provides a direct measure of chromatin states involved in gene expression, such as the occupation of promoter and enhancer sequences by transcription factors and other proteins. We assayed chromatin accessibility using ATAC sequencing (Buenrostro et al. 2013) in leaf mesophyll protoplast nuclei of hexaploid Paragon wheat and diploid *Ae. tauschii.* Comparison of ATAC peak lengths on chromosomes 3DL and 3L identified major differences. Chromatin in promoter and 5’UTR regions of genes in the diploid context had more accessible chromatin, both in terms of peak numbers and peak lengths, than in the hexaploid context (Fig. 6A, B, E). This restriction of chromatin access in the hexaploid context extended across the chromosome arm and encompassed genes with both proportionately reduced and differential expression (Fig. 6F). Proportionately reduced gene expression on 3DL was correlated with reduced chromatin accessibility in hexaploid Paragon compared to diploid *Ae. tauschii* (Fig. 6B), and nearly all DEGs also displayed reduced chromatin accessibility in Paragon (Fig. 7). Interestingly, although the bulk of non-genic regions of 3DL and 3L had relatively similar extents of inaccessible chromatin compared to genic regions, a small proportion of annotated repeats, predominantly simple sequence repeats, had regions of accessible chromatin (Fig. 5A, B). Accessible chromatin was also found in class I and class II repeats, and these might be associated with repeat activities. The relatively high levels of chromatin accessibility in SSRs may reflect possible altered nucleosome binding dynamics to these atypical sequences, as Alu repeats influence nucleosome spacing in human cells (Tanaka et al. 2010). In human cancer cells a small set of SSRs had increased chromatin accessibility during differentiation and this was linked to the regulation of specific genes (Gomez et al. 2016). We did not observe a pattern of SSRs adjacent to genes that had differential expression or chromatin accessibility; most SSR with accessible chromatin were scattered over Mb distances from genes on 3DL and 3L (data not shown), which did not support a clear role in regulation of adjacent genes. Overall, the reduction in chromatin accessibility was much less in non-gene space regions of 3DL compared to gene space (Fig. 6A, B), showing that hexaploidization leads to preferential reduction in chromatin accessibility in genes compared to intergenic regions.

Differences in chromatin accessibility, as measured by ATAC, are thought to be due to passive competition for DNA between nucleosomes and transcription factors, chromatin remodelling and architectural proteins (Klemm et al. 2019). It is possible that in the hexaploid context reduced chromatin access across genic regions of 3DL may be due to altered competition for DNA between increased nucleosome formation or reduced transcription factor levels, or a combination of both. The overall similarities in chromatin accessibility in intergenic regions (Fig. 5B) of both 3L ands 3DL may be due to higher-order nucleosome packaging in heterochromatin, which is characteristic of intergenic DNA in larger grass genomes (Baker et al. 2015). It is conceivable that the introduction of a divergent set of gene regulatory proteins from the 4Gb *Ae. tauschii* genome into a 12Gb tetraploid nucleus to form an allohexaploid wheat genome leads to altered dynamics between the new set of transcription factors and nucleosomes, thus altering chromatin accessibility. Models of homoeolog expression patterns in allopolyploids have incorporated varying affinities and concentrations of transcription factors for gene regulatory sequences in an allopolyploid (Hu and Wendel 2018). This showed a strong effect of inter-genome interactions on altered gene expression. However, other mechanisms may also contribute to establishing gene expression patterns in allopolyploid genomes. Recently, the conformation of cotton chromosomes was shown to be strongly affected by polyploidy, which altered Topologically Associating Domain (TAD) boundaries and chromatin states (Wang et al. 2018). Similar changes may occur on wheat upon polyploidization. In interphase nuclei of grasses such as hexaploid wheat, chromosomes adopt the Rabl configuration, with centromeres clustered at one side of the nucleus and telomeres at the other, such that homoeologous chromosome arms are aligned (Abranches et al. 1998). There was no evidence at this optical scale for clustering of transcriptional sites. DNA proximity assays have shown larger-scale interactions expected from the Rabl configuration in diploid barley (Mascher et al. 2017), but these have not yet been carried out in allopolyploid wheat at the resolution required to detect any proximity of smaller genomic segments such as TADs or putative homoeologous interacting domains. Our identification of concerted changes in chromatin accessibility in a pair of homologous chromosome arms in the diploid and hexaploid genome context has started to identify mechanisms that establish and maintain balanced and differential gene expression patterns in polyploid genomes such as wheat. It will be important to assess gene expression and chromatin accessibility across the complete genomes of newly formed allopolyploid wheat lines to identify mechanisms that rapidly establish and maintain the extensive differences in gene expression caused by alllopolyploidy in wheat, and perhaps in other polyploid genomes.

## Methods

### Plant Materials

The Paragon elite hexaploid spring wheat *(Triticum aestivum*, AABBDD) variety was used as it is a commonly used experimental line with a sequenced genome and extensive genetic resources. The diploid progenitor species *Aegiiops tauschii* (DD), accession AL8/78 has a sequenced genome was used for most experiments. Two divergent *Ae. tauschii* lines, Clae23 and ENT336 were also used for comparative transcriptomics. All plants were grow in a glasshouse with supplementary lighting (12-24 °C, 16/8 light/day). Young leaf root tissue and whole seedlings, and developing grain samples were collected for RNA extraction. All tissue samples were immediately frozen in liquid nitrogen and stored at −80 °C.

### Chromosome 3DL Arm Assembly and Annotation

A full description is provided in Additional File 1. Briefly, a set of BAC scaffolds of flow-sorted chromosome 3DL (Lu et al. 2018) was extended and scaffolded using wheat PacBio assemblies from Triticum 3.1 (Zimin et al. 2017) as templates. These scaffolds were further extended using Fosill long mate-pair reads (Lu et al. 2018). The chromosome 3DL pseudomolecule was created by mapping the resulting 504 scaffolds to IWGSC 3D (International Wheat Genome Sequencing Consortium (IWGSC) et al. 2018) with MUMmer v3.23. The scaffolds were localised and assigned to a specific order and strand and linked using scaffolder v0.5.0. Order discrepancies were manually corrected and one hundred Ns were placed between neighbouring scaffolds to mark the sequence gap. Chromosome 3DL genes were predicted by *ab initio methods*, using EST and *de novo* transcript assemblies. Predicted gene models were curated manually using Integrated Genome Viewer (IGV) and given a confidence score using RNA evidence, protein alignments (Additional File 2: Table S8) and *ab initio* predictions. Pseudogenes were annotated and identified as those genes with predicted exon-intron structures conforming to the GT-AG intron rule, but with no consensus or translatable CDS. Repeats were identified using RepeatMasker and sequence comparisons with RepBase. Gene ontology searches used blastx against the NCBI NR protein database and the the top 20 hits were used for GO-mapping. GO terms were assigned to genes using BLAST2GO.

### Gene synteny

*Ae. tauschii* AL8/78 genome pseudomolecules and their gene annotations were from (Luo et al. 2017). An additional 1,266 additional genes were identified on chromosome 3L using wheat 3DL gene models and *Ae. tauschii* RNAseq and transcript assemblies. A total of 4,121 genes were assigned to chromosome 3L. Assemblies and annotations of the related grasses *Brachypodium distachyon, Hordeum vulgare, Oryza sativa* Japonica, and *Sorghum bicolor* were downloaded from Ensembl Plants (version 38). Similarity searches between wheat and these species were performed using BLAST+ (v2.6.0; parameter: -evalue 1e-10 -outfmt 6 -num_alignments 5) (Camacho et al. 2009), and gene synteny and collinearity were detected using MCScanX with default settings (Wang et al. 2012), and plotted using VGSC (v1.1) (Xu et al. 2016).

### Transcription Analyses

A full description is provided in Additional File 1. Triplicated samples of Paragon wheat and *Ae. tauschii* AL8/78 were collected from leaves and roots of 14 day old greenhouse grown plants, 4 day old seedlings germinated on filter paper, and 10 DAP and 27 DAP developing grains (Additional File 2: Table S2), frozen in liquid nitrogen and stored at −80 °C. Total RNA was extracted as described (Oñate-Sánchez and Vicente-Carbajosa 2008). Illumina TruSeq mRNA libraries were constructed according to manufacturer’s protocol. All sequencing was carried out on an lllumina HiSeq 2500, with 100 bp paired-end read metric, TruSeq SBS V3 Sequencing kit and version 1.12.4.2 RTA. FASTQ files were generated and demultiplexed according to library-specific indices by CASAVA (v. 1.8.2, lllumina). Reads were mapped to the complete Triticum3.1 genome assembly with 3DL assemblies replaced by 2,703 chromosome 3DL scaffolds. HISAT2 (v2.1.0) (Kim et al. 2015) was used to create the genome indices and map the trimmed RNAseq reads to each genome. Expression differences between Paragon and AL8/78 used the HISAT2-StringTie pipeline (Pertea et al. 2016) to compute Transcripts Per Million (TPM) values. Absolute quantitative RT-PCR was used to validate TPM value comparisons between the diploid and hexaploid chromosome arms. Fourteen genes with 1:1:1 A:B:D homoeologs and balanced expression were identified and two pairs of primers for RT-PCR were designed. One pair was designed to amplify all three gene copies in hexaploid wheat and the single copy in diploid *Ae. tauschii*, and a second pair was designed to specifically amplify only the DD genome copy in hexaploid wheat and diploid *Ae. tauschii.* Triplicate samples of 50,000 leaf protoplasts from Paragon wheat and *Ae. tauschii* AL8/78 from young leaves were collected and RNA extracted. Quantitative PCR was performed on cDNA using a LightCycler 480 System (Roche), and a standard curve of 8-80m molecules of the 5,340-bp plasmid pETnT was used to estimate absolute transcript levels.

### Bisulphite Sequencing

A full description is provided in Additional File 1. Triplicated DNA samples from *Ae. tauschii* AL8/78 and Paragon wheat 14 day old leaves were extracted for analysis. For Paragon wheat, whole genome bisulphite sequencing was carried out, while gene capture with Agilent SureSelect Target Enrichment was used for *Ae. tauschii* chromosome 3L predicted genes. Bisulfite treatment of triplicated samples used the Zymo Research EZ DNA Methylation-Gold Kit, standard lllumina library preparation and sequencing using a Hiseq 4000 (2 × 150 bp reads) for *Ae. tauschii* samples and a Hiseq 2500 (2 × 250 bp reads) for Paragon wheat samples. Bisulfite converted Paragon paired end sequences were aligned to Paragon genome assemblies using Bismark (version 0.18.1) (Krueger and Andrews 2011) and chromosome 3DL methylation status was identified by alignment of 13,046,879 bp of Paragon methyl-sequences with the Chinese Spring 3DL pseudomolecule. For *Ae. tauschii*, 19,519,314 bp of methyl-sequences across 4,130 sequences (18,975,440 bp of unique sequence) were identified as part of chromosome 3L. The methylation status of each cytosine residue across the sequences was identified using the Bismark methylation extractor tool and the percentage of reads methylated per cytosine residue across the wheat 3DL and *Ae. tauschii* sequences were calculated.

### ATAC sequencing

A protocol adapted for wheat and *Ae. tauschii* leaf nuclei is described in Additional File 1. Nuclei were isolated rapidly from triplicated leaf protoplast samples and subjected to transposition using Nextera transposase (lllumina). Naked DNA was used as a control. Amplified tagmented DNA was sequenced on a HiSeqX lllumina sequencer with 150 bp paired-end reads (Novagene, Shenzhen). ATAC-seq reads matching mitochondrial and chloroplast genomes were identified and removed using the *Triticum aestivum* chloroplast genome (GenBank accession No. NC_002762), *Triticum aestivum* mitochondrial genome (GenBank accession No. AP008982) and the *Ae. tauschii* chloroplast genome (GenBank accession No. NC_022133). The wheat mitochondrial genome was used to identify *Ae. tauschii* mitochondrial reads. After filtering, paired reads with high mapping quality (MAPQ score >10, qualified reads) were identified using SAMtools (Li et al. 2009). Duplicate reads was removed using the Picard tools MarkDuplicates program (http://broadinstitute.github.io/picard/. All reads aligning to the forward strand were offset by +4bp, and all reads aligning to the reverse complement strand were offset by −5 bp (Adey et al. 2010). ATAC-Seq peak regions of each sample were called using MACS2 (v2.1.2_dev) (Zhang et al. 2008) and ATAC-Seq peaks for which the distance between proximal ends was less than 10 base pairs were merged.

### Availability of data and material

The chr3DL pseudomolecule assembly and unlocalised scaffolds, RNAseq and ATAC-seq data were submitted to European Nucleotide Archive (ENA). chr3DL assembly and annotation – ERZ795128 under PRJEB23358; Paragon RNAseq – PRJEB29855; *Ae tauschii* AL8/78 RNA-seq – PRJEB23317 as described in the *Ae. tauschii* genome paper (Luo et al. 2017); *Ae tauschii* Clae 23 leaf RNA-seq – PRJEB29859; *Ae tauschii* ENT336 leaf RNA-seq – PRJEB29860; Paragon ATAC-seq – PRJEB29868; AL8/78 ATAC-seq – PRJEB29869. Gene methylation data for *Ae tauschii* AL8/78 3L and Paragon wheat 3DL is in PRJEB31186.

## Acknowledgements

We are grateful to project IOS1238231 of the US National Science Foundation for providing access to unpublished sequence of the *Ae. tauschii* genome. We are also grateful to Dr Tarang Mehta from the Earlham Institute for advice on ATAC protocols. We thank the sequencing teams at The Earlham Institute, Cold Spring Harbor Laboratory and University of Liverpool CGR for their expert and timely generation of sequencing data.

## Authors’ contributions

F-HL and MWB conceived and managed the research. F-HL conducted bioinformatics analyses. NMcK and F-HL carried out laboratory work. L-JG performed DNA methylation analyses. AH and M-CL provided material and advice prior to publication. MWB and F-HL wrote the paper with contributions from L-JG and NMcK. All authors have seen and approved the manuscript.

## Additional Files

**Additional File 1** Supplementary Methods

**Additional File 2** Supplementary Figures and Tables. **Figure S1** Expression profiles of syntenic genes of *Triticum aestivum* Paragon 3DL and *Aegilops tauschii* AL8/78 3L in in five tissues. **Figure S2** Comparison of expression levels of between *Aegilops tauschii* varieties AL8/78, Clae23 and ENT336. **Figure S3** Illustration of gene structure differences among 106 conserved DEGs between wheat and *Ae. tauschii.* **Figure S4** Size distribution of ATAC-seq fragment lengths between *Triticum aestivum* Paragon and *Aegilops tauschii* AL8/78. **Figure S5** Normalised read enrichments for four classes of chromosome states in *Triticum aestivum* Paragon and *Aegilops t*auschii AL8/78. **Table S1.** Summary of repetitive elements of *Triticum aestivum* Chinese Spring 3DL and *Aegilops tauschii* AL8/78 3L. **Table S2** Number of RNAseq read pairs after trimming in 5 sampled tissues of *Triticum aestivum* Paragon and *Aegilops tauschii* AL8/78. **Table S3** Number of differential expressed genes in 5 tissues between *Triticum aestivum* Paragon and *Aegilops tauschii* AL8/78. **Table S4** Mapping statistics after alignment of bisulfite treated *Triticum aestivum* Paragon and *Ae. tauschii* AL8/78 samples. **Table S5** Number of regions for methylation analysis in *Triticum aestivum* Paragon 3DL and *Aegilops tauschii* AL8/78 3L. **Table S6** Summary of ATAC peaks and covered genes in different chromosome states of Paragon wheat and *Ae. tauschii* AL8/78. **Table S7** Number of syntenic genes covered by ATAC-seq peaks. **Table S8** Protein sequences used as evidence for exons in wheat chromosome 3DL annotation.

**Additional File 3** Statistics of 504 BAC scaffolds and the alignments to IWGSC v0.4 wheat 3DL and *Ae. tauschii* 3L

**Additional File 4** Syntenic genes and RNAseq levels expressed as TPMs

**Additional File 5** Primers used for quantitative real-time PCR

**Additional File 6** Gene Ontology annotations of **173** differential expressed genes in **27** DAP developing grain

