## Additional File 1 for "Reduced chromatin accessibility underlies gene expression differences in homologous chromosome arms of hexaploid wheat and diploid *Aegilops tauschii*"

### Additional File 1. Supplementary Methods

#### Wheat chromosome 3DL Assembly and Annotation

A total of 2,703 PacBio scaffolds from Chinese Spring chromosome 3DL were retrieved (Zimin et al. 2017) and overlapped and extended to 1,751 scaffolds using FosiIl mate-paired reads as described previously (Lu et al. 2018). These 1,751 PacBio scaffolds were then integrated with our 3DL BAC scaffolds (International Wheat Genome Sequencing Consortium (IWGSC) et al. 2018; Lu et al. 2018) to generate a final set of 524 super-scaffolds using nucmer (default setting) and delta-filter (-i 99 -l 500) in MUMmer v3.23 (Kurtz et al. 2004) to identify joins between BAC and PacBio scaffolds, and further assembled using scaffolder v0.5.0 software (Barton and Barton 2012). BAC scaffolds were aligned to 524 templates using nucmer (default setting) and further filtered using delta-filter (-i 99 -r -q -l 500), and then were further merged to form 504 scaffolds by a custom Perl script designed to merge BACs for each template scaffolds using quickmerge v0.2 (-hco 5.0 -c 1.5 -l 1000 -ml 500) software (Chakraborty et al. 2016). A 3DL pseudomolecule was made by mapping 504 scaffolds to the IWGSC chr3D pseudomolecule (International Wheat Genome Sequencing Consortium (IWGSC) et al. 2018) using MUMmer v3.23. All scaffolds were localised and assigned to a specific order and strand and then linked using scaffolder v0.5.0. Order discrepancies were manually corrected. One hundred Ns were placed between two neighbour scaffolds to mark the sequence gap.

Repetitive sequences of the 3DL pseudomolecule were identified using RepeatMasker (v4.0.7) with the submodule Tandem Repeat Finder (v4.09) (Benson 1999). Sequence comparisons were performed using the alignment software RMBlast (v2.2.28). A database of repetitive DNA (109,726 sequences) were collected from Repbase RepBase (v20170127; 45,447 sequences) (Bao et al. 2015), PGSB-REdat(v9.3p; 61,730 sequences) (Nussbaumer et al. 2013) and 2,549 repetitive substrings (4,671,512 bp) with at least 10-time appearance that were discovered by RepeatScout (v1.0.5) (Price et al. 2005). Those 2,549 repetitive substrings were classified by PASTE Classifier in REPET package (v2.5) (Hoede et al. 2014) using the REPbase databases (REPET edition v20.05 with 38,777 nucleotide sequences and 24,192 amino acid sequences).

Genes were identified *ab initio* on the chromosome 3DL pseudomolecule using Augustus (v3.0.3) (Stanke et al. 2006) trained for hexaploid wheat. A training set was made from 7,264 CDS sequences from wheat 3B (Choulet et al. 2014) mapped to 3DL pseudomolecule using the PASA pipeline (version r20140417; parameters: --runpasa -a gmap,blat) (Haas et al. 2008). Those genes, overlapped with each other or with similarity >=70% or bad genes (number of bases cannot be divided by three), are removed and not used for following analysis. It generated a total of 787 gene models, of which 687 were used to train Augustus to generate wheat-specific prediction parameters, and the remaining were used for testing the precision of these trained prediction parameters. To clearly show and refine the wheat exon borders covered by Paragon RNAseq reads, a two-step RNAseq mapping strategy was employed using Gsnap2Augustus (https://github.com/lufuhao/Gsnap2Augustus). Firstly, RNAseq reads were mapped to the 3DL pseudomolecule using GSNAP (version: 2017-12-29; parameter: --nofails --expand-offsets=1 --novelsplicing=1 -B 3 --localsplicedist=10000 --npaths=30 --format=sam) (Wu and Nacu 2010). This generated a preliminary *ab initio* prediction of BAM intron hints. An exon-exon database was created using the script 'intron2exex.pl' included in Augustus. Secondly, RNAseq reads were mapped to the exon-exon database and the coordinates were calibrated back to the 3DL pseudomolecule using 'samMap.pl' included in Augustus. Precise BAM intron hints were generated by bam2hints included in Augustus. Finally, Augustus calculated *ab initio* predictions based on BAM intron hints.

EST evidence for gene predictions were generated using Exonerate (v2.4.0; parameter: --model est2genome --percent 70 --score 100 --showvulgar yes --bestn 10 --minintron 20 --softmaskquery no --softmasktarget no --showalignment no --showtargetgff yes --geneseed 250) (Slater and Birney 2005) using 3 EST datasets: *de novo* assembly of Paragon RNAseq reads using the Trinity assembler (v2.0.6; parameter: --genome_guided_max_intron 10000) (Grabherr et al. 2011); the Triticeae Full-Length CDS Database (TriFLDB) (Mochida et al. 2009); and 1,551,792 *Triticum aestivum* ESTs downloaded from NCBI. Protein evidence was produced by mapping protein sequences from 10 species (Additional File 2: Table S8) to the 3DL pseudomolecule using GenomeThreader (v1.6.2; parameter: -gff3out yes -skipalignmentout yes) (Gremme et al. 2005). EVidenceModuler (EVM; v20120625) (Haas et al. 2008) was used to combine *ab initio* gene predictions, EST and protein alignments into weighted (1:5:10) consensus gene structures. The pipeline and converters are available at https://github.com/lufuhao/AutoEVM.

EVM gene models were manually curated using the Integrative Genomics Viewer (IGV; v2.3.60) (Thorvaldsdóttir et al. 2013) and given a confidence score (0-5), in which evidence from RNAseq peaks, *ab initio* predictions, protein alignments, *de novo* EST alignments and NCBI EST alignments separately accounted for a single confidence value. Pseudogenes were annotated as genes with good exon-intron structures that conformed to the GT-AG intron rule, but had no consensus/translatable CDS. Finally, manually curated gene modules were transferred to the chr3DL pseudomolecule using RATT (in PAGIT v1) (Otto et al. 2011) and some remaining modules using custom scripts (https://github.com/lufuhao/ExonerateTransferAnnotation), which employed Exonerate to map transcripts and CDS sequences separately and then integrated them together into final gene models.

Similarity searches were carried out using blastx in the BLAST+ toolkit (v2.6.0; parameter: -evalue 1e-6 -outfmt 5 -show_gis -num_alignments 20 -max_hsps 20) against the NCBI NR (v20171024) protein database. The best 20 hits with E-value 1E-6 for each sequence were retained and used for GO-mapping. GO terms (version 07-Jan-2017) associated with these candidate sequences were assigned to each gene using BLAST2GO (v2.5; parameter: -v -annot -dat -img -ips ipsr -annex -goslim) (Götz et al. 2008). The web plotting tool WEGO (Ye et al. 2006) was used to draw GO annotations.

#### Gene Expression Analyses

RNA samples from Paragon and *Ae. tauschii* lines AL8/78, Clae23 and ENT336 were extracted as described by Oñate-Sánchez and Vicente-Carbajosa (Oñate-Sánchez and Vicente-Carbajosa 2008). RNA quality was assessed using a Qubit fluorometer (Invitrogen) and 2100 Bioanalyzer (Agilent Technology) in the Earlham Institute (Norwich, UK). Illumina TruSeq mRNA libraries were constructed using the Illumina TruSeq RNA Sample preparation guide v2 (Illumina Inc.) in accordance with the manufacturer’s protocol. One µg of total RNA used to purified mRNA using two rounds of poly-T oligonucleotide purification attached magnetic beads. During the second elution of poly-A RNA, the RNA was fragmented and primed for cDNA synthesis. cDNA synthesis was carried out using SuperScript II Reverse Transcriptase (Invitrogen) and random primers. Second strand cDNA synthesis was carried out and the DNA was subjected to end repair, “A” tailing and ligation. cDNA templates were enriched by 15 cycles of PCR as per manufacturer's instructions. The amplified library was quantified using a Bioanalyzer DNA 100 Chip. The library was normalised to 10 nM for generation of sequence clusters on a sequencing flow-cell on the Illumina c-Bot instrument. Sequencing library cluster generation was carried out on a paired-end flow cell on the Illumina cBot according to the manufacturer's instructions. All sequencing was carried out on an Illumina HiSeq 2500, with 100 bp paired-end read metric, TruSeq SBS V3 Sequencing kit and version 1.12.4.2 RTA. FASTQ files were generated and demultiplexed according to library-specific indices by CASAVA (v. 1.8.2, Illumina). Adaptor sequences were trimmed using CutAdapt v 1.6 (Martin 2011) and low quality sequences were removed by Trimmomatic v0.30 (Bolger et al. 2014) with parameter ILLUMINACLIP:2:30:10 HEADCROP:10 LEADING:3 TRAILING:3 SLIDINGWINDOW:4:15 MINLEN:36 (Additional File 2: Table S2).

To estimate gene expression differences between Paragon and *Ae. tauschii*, the HISAT2-StringTie pipeline (Pertea et al. 2016) was used to compute Transcripts Per Million (TPM) values. Transcripts were mapped to the Triticum3.1 wheat assembly (Zimin et al. 2017) with the 2,703 PacBio-based 3DL scaffolds replaced by the 3DL pseudomolecule, and to the *Ae. tauschii* AL8/78 assembly (Luo ref).. HISAT2 (v2.1.0) (Kim et al. 2015) was used to create genome indices and map the trimmed RNAseq reads to each genome. A two-step StringTie (v1.3.3b) (Pertea et al. 2016) strategy was used to measure transcript abundance. For each RNA-Seq sample, StringTie was used to assemble the read alignments; and then a non-redundant set of transcripts observed in all the RNA-Seq samples assembled previously was generated using the stringtie --merge mode; finally the second run was performed with the merged transcript models, for each RNA-Seq sample, using the -B/-b and -e options in order to estimate transcript abundances and generate read coverage tables.

Absolute quantitative RT-PCR was employed to validate the use of TPM values for comparing TPM values between species. Fourteen genes exhibiting balanced expression that had single copy AA and BB homoeologs in the TGACv1 wheat genome were selected for primer design for Q-RT-PCR using HomoeologPrimer (https://github.com/lufuhao/HomoeologPrimer). Two pairs of primers for each gene was designed, one pair to amplify all AA+BB+DD transcripts in hexaploid wheat and the DD transcript in diploid *Ae. tauschii*, and other pair to specifically amplify DD transcripts in the hexaploid and the diploid genomes. For each species, 50,000 protoplasts from young leaves were collected in triplicates, total RNA made using the Qiagen

RNeasy Plant Mini Kit (Cat No. 74904). Up to 1 µg total RNA was was used to generate cDNAs using Qiagen QuantiTect Reverse Transcription Kit (Cat No. 205311), which yielded 20 µl of cDNAs (33 ng/ul for Paragon and 26 ng/ul for AL8/78). Quantitative PCR was performed on the LightCycler 480 System (Roche). A dilution curve of 8 - 80m copies of the 5,340-bp plasmid, pETnT was used to to establish a standard curve of absolute molecule numbers. The 20-µl PCR profile was set as follows: 10 µl of LightCycler® 480 SYBR Green I Master, 1 µl of forward primer, 1 µl reverse primer, 0.3 1 µl cDNA, 7.7 µl of $H_2O$; The LightCycler 480 instrument was run as follows: 10 min of pre-incubation at 95 ℃; 45 cycles amplification: 10s at 95 ℃, 10s at 72 ℃ and 10s at 72 ℃ with single acquisition; Melting Curve and cooling was set as instructed in the SYBR Green I Master protocol.

#### Bisulphite Sequencing

Leaf material was harvested from triplicated samples DNA was extracted using the Qiagen DNeasy Plant Mini Kit. Samples from *Ae. tauschii* were processed at the CGR (University of Liverpool, UK) using the Agilent SureSelect capture probe sets for targeted gene enrichment followed by bisulfite treatment using the Zymo Research EZ DNA Methylation-Gold Kit, standard illumina library preparation and sequencing using the Hiseq 4000 (2 x 150 bp reads). Triplicated samples from hexaploid Paragon wheat were pooled and processed to generate a whole genome bisulfite treated sequencing library using the Zymo Research EZ DNA Methylation-Gold Kit and sequencing was carried out on a Hiseq 2500 at Cold Spring Harbor Laboratories, USA (2 x 250 bp reads). Bisulfite-converted Paragon and *Ae. tauschii* paired end sequences were aligned to the full Paragon genome assembly or *Ae. tauschii* 3L assemblies using Bismark (version 0.18.1) (Krueger and Andrews 2011). Duplicate sequencing reads were then filtered using Picard tools. The Bismark methylation extractor tool was then used to identify the methylation status at each cytosine residue across the sequencing reads. A custom Perl script was then used to calculate the % of reads methylated per cytosine residue across the reference sequence. To map Paragon methylation data to the 3DL pseudomolecule, methylation data was aligned to the Paragon genome assembly (https://opendata.earlham.ac.uk/opendata/data/Triticum_aestivum/) using Nucmer (Kurtz et al. 2004) and then the methylation coordinates were transferred to the Chinese Spring 3DL pseudomolecule. The longest contiguous alignments between the sequences with identities >98% and lengths ≥500 bp were identified and the methylation status of 3DL genic DNA was identified in the pseudomolecule. This yielded a space of 13,046,879 bp across 3,541 sequences that were identified as part of the 3DL pseudomolecule. For *Ae. tauschii*, genic sequences in chromosome 3L spanned 19,519,314 bp across 4,130 sequences (18,975,440 bp of unique sequence).

Gene sequences for probe design for targeted gene enrichment included 80,562,496 bp from *Ae. tauschii* chromosome 3L. This included 2000 bp upstream of each gene (from the start codon), the gene body including predicted introns, and 500 bp downstream of the termination codon. Agilent SureSelect Target Enrichment probes contained 120 bp probes tiled across the design space at 40 bp intervals. A total of 227,969 probes covering a potential 27,356,280 bp were submitted for synthesis using the Agilent SureDesign capture design website.

#### ATAC Sequencing

Sterilised seeds were grown on moist filter paper in petri dishes at 25°C for 11-12 days. Leaf tissue (1 - 2 g) was cut into 2 - 3 cm lengths in a sterile petri dish containing 10 ml 0.6 M Mannitol, 3% Cellulase RS (Duchefa), 1% Macerozyme R10 (Duchefa), 10mM MES, pH5.7, 1mM $CaCl_2$, 5mM *ß*-Mercaptoethanol, 0.1% BSA, 50 ppm ampicillin. Leaf material was finely chopped to 1 - 2 mm segments and vacuum infiltrated at 20 mm Hg for 20 mins. Protoplasts were released by shaking (50 rpm) at 25 °C for 4-4.5 hour in the dark. The lysate was filtered through a 100 µm Cell Strainer (Falcon) into a 50 ml Falcon tube, and the digested leaf material was washed three times with 6 ml suspension buffer (0.6 M Mannitol, 20mM KCl, 4mM MES, pH5.7 with KOH). The filtrate was re-filtered through a 70 µm Cell Strainer (Falcon) and centrifuged at 70g for 15 min at 12 °C. The protoplast pellet was gently resuspended in 2 ml of suspension buffer and layered onto a Percoll gradient (2.25 ml Percoll (Sigma) and 5.25 ml Suspension buffer) in a 15ml Falcon tube. The gradient was centrifuged at 3000 rpm for 15 min, and the upper phase was discarded and the protoplast layer (cloudy light green phase) was collected into new 15 ml Falcon tube. This was washed in 9 ml suspension buffer, centrifuged at 70 g for 10 min, and the pellet resuspended in 2 ml suspension buffer. Protoplast yields and integrity were tested using Evans Blue (400 mg/l in 0.5M mannitol) and a haemocytometer. This yielded more that 200,000 viable high quality protoplasts.

Nuclei were prepared by pelleting approximately 200,000 protoplasts at 70g for 10 min at 12 °C and resuspending in 6 ml MEB buffer containing 0.1% Triton X-100 (1.0M 2-methyl-2,4-pentanediol (Aldrich),10 mM PIPES-KOH, 10 mM $MgCl_2$, 0.1% Triton X-100, 2%

polyvinylpyrrolidone (PVP-10 Sigma), 10mM sodium metabisulfite, 5 mM mercaptoethanol, pH 6.0) and mixed by gentle rotation at 4°C for 5 mins. The crude nuclear prep was centrifuged at 650g for 5 mins at 4°C and resuspended in 3 ml MPDB buffer containing 0.1% Triton X-100 buffer (0.375M 2-methyl-2,4-pentanediol (Aldrich), 7.5 mM PIPES-KOH, 7.5 mM $MgCl_2$, 0.1% Triton X-100, 7.5 mM sodium metabisulfite, 5 mM mercaptoethanol, pH 7.0). The suspended nuclei were layered onto a 37.5% Percoll gradient (3.75ml Percoll,(Sigma) and 6.25ml MPDB 0.1% Triton X-100 buffer) in a 15ml Falcon tube and centrifuged at 1000g for 10 mins at 4°C. The purified nuclear pellet was resuspended in 7.5 ml MPDB 0.1% Triton X-100 buffer, centrifuged at 650g for 5 mins at 4°C, and resuspended in 1-2 ml MPDB 0.1% Triton X-100 buffer. The yield and integrity of purified nuclei was assessed using a haemocytometer and Methylene blue staining. Typical yields were 25-30% of starting protoplast numbers. Nuclear preparation should be completed within 30 mins of protoplast lysis.

For ATAC reactions on nuclei, reagents were from the Illumina Nextera DNA Library Prep Kit FC-121-1030/15028212 (24 samples). For a single reaction 50,000 intact isolated nuclei were pelleted in a microfuge at 650g 5 mins 4 °C. The nuclear pellet was resuspended in 25 µl 2x Tagment DNA Buffer. 22.5 µl or 20 µl nuclease free water and 2.5 µl or 5.0 µl Tagment DNA Enzyme 1 was added and the reaction transferred to a 0.2ml PCR tube and incubated in thermal cycler block pre-warmed to 37°C for 30 mins, with gentle mixing by hand every 5 mins. Immediately following the transposition reaction, DNA was purified using a Qiagen PCR Purification MinElute kit. Elute DNA with 12 µl Elution Buffer and resuspended in 10 µl. Tagmented DNA can be stored at -20°C at this stage.

Amplification reactions were set up in a PCR tube:

| | |
|---|---|
| Tagmented DNA | 10 µl |
| Nuclease Free Water | 10 µl |
| 25uM Customized Universal P7 Primer | 2.5 µl |
| 25uM Customized Barcoded P5 Primer | 2.5 µl |
| NEB High Fidelity 2x PCR Master Mix | 25 µl |

PCR cycles:

| | | |
|---|---|---|
| 1 Cycle | 5 Mins | 72°C |
| | 30 Secs | 98°C |
| 11 Cycles | 10 Secs | 98°C |

| | | |
|---|---|---|
| | 30 Secs | 63°C |
| | 1 Min | 72°C |
| Hold | | 4°C |

Amplified DNA was purified using a Qiagen PCR Purification MinElute kit. DNA was eluted with 22 µl Elution Buffer. An additional clean up step to removed excess primers used 20 µl Ampure XP Beads (Beckman Coulter A63880) and purified DNA was recovered in 25 µl 0.1 x TE. DNA was quantified using a Qubit Fluorometer and a HS DNA kit. The size distributions of tagmented amplified DNA was assessed using MultiNA MCE-202 Bioanalyser (Shimadzu) with a DNA 1000 Kit, or Tapestation Screen tape Kit D1000, or the Agilent Bioanalyser HSDNA chip.

To exclude ATAC-seq reads from mitochondrial and chloroplast genomes, which arise from contamination of plant nuclei preparations, we included the *Triticum aestivum* chloroplast genome (GenBank accession No. NC_002762), *Triticum aestivum* mitochondrial genome (GenBank accession No. AP008982) and *Ae. tauschii* chloroplast genome (GenBank accession No. NC_022133) in the reference genome. As the *Ae. tauschii* mitochondrial genome has not yet been sequenced that from wheat was used. Trim_Galore (version 0.5.0; https://github.com/FelixKrueger/TrimGalore) was used to remove Nextera adaptor sequences and Trimmomatic (Bolger et al. 2014) were used to filter out those short (<70 bp) reads. The resulting clean reads were aligned to the reference sequences using Bowtie (v1.2.2; Options: -X 2000 --fr -m 1) (Langmead et al. 2009). After filtering reads matching mitochondrial and chloroplast genomes (which also removed reads matching chloroplast and mitochondrial insertions in the nuclear genome), we included paired reads with high mapping quality (MAPQ score >10, qualified reads) through SAMtools (Li et al. 2009) for further analysis. Duplicate reads was removed using Picard tools MarkDuplicates (http://broadinstitute.github.io/picard/). All reads aligning to the forward strand were offset by +4bp , and all reads aligning to the reverse complement strand were offset -5 bp (Adey et al. 2010). ATAC-Seq peak regions of each sample were called using MACS2 (v2.1.2_dev) (Zhang et al. 2008) with parameters --nomodel --shift -37 --extsize 73. To generate a consensus set of unique peaks, we next merged ATAC-Seq peaks for which the distance between proximal ends was less than 10 base pairs.
