## Additional File 2 for "Reduced chromatin accessibility underlies gene expression differences in homologous chromosome arms of hexaploid wheat and diploid *Aegilops tauschii*"

### Additional File 2. Supplementary Figures and Tables

**A**

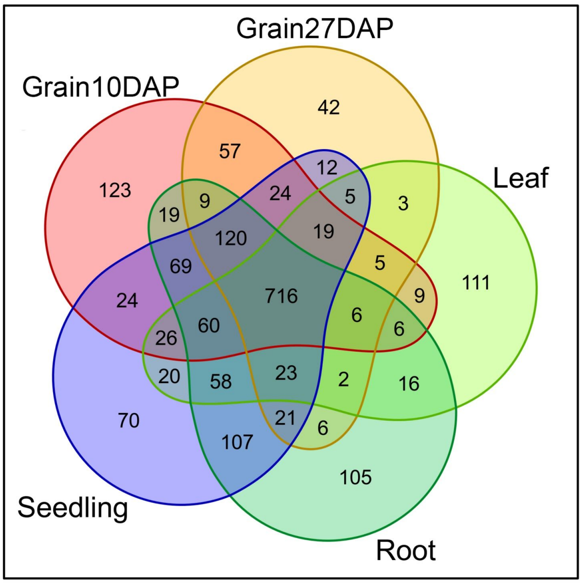


**B**

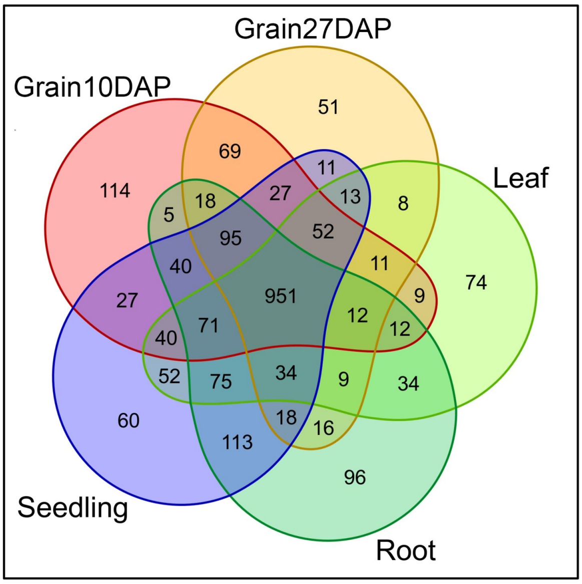


**Figure S1 Expression profiles of syntenic genes of *Triticum aestivum* Paragon 3DL and *Aegilops tauschii* AL8/78 3L in five tissues.**

Syntenic genes expressed with TPM≥1 from wheat 3DL (1,893 genes, **A**) and *Ae. tauschii* chr3L (2,217 genes, **B**) among leaf, root, seedling, 10 DAP (Days After Pollination) and 27 DAP developing grain.

A

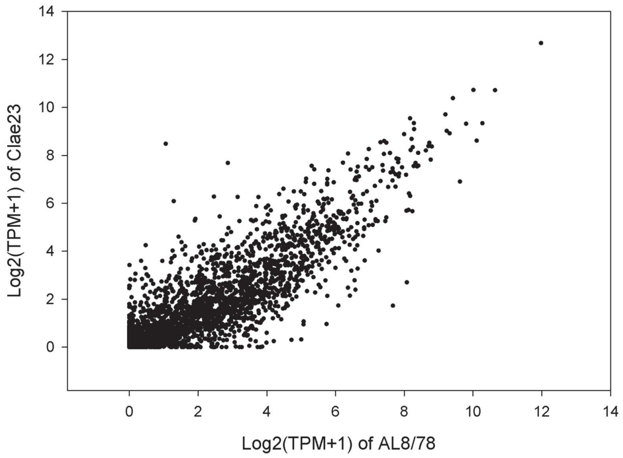


B

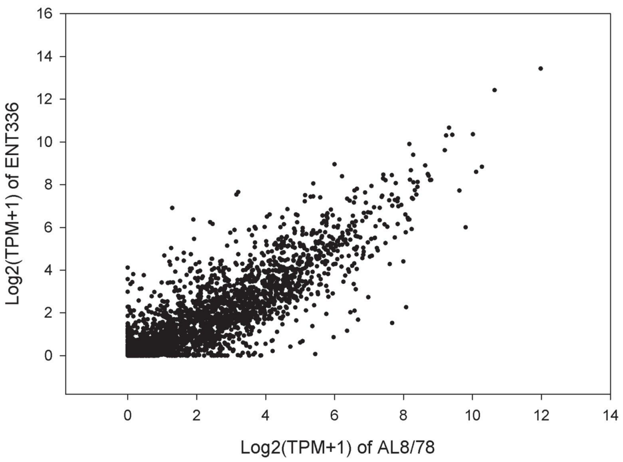


**Figure S2 Comparison of expression levels of between *Aegilops tauschii* varieties AL8/78, Clae23 and ENT336.**

To identify conserved differentially expressed genes (DEGs) between *Triticum aestivum* Paragon and *Aegilops tauschii* AL8/78, RNAseq data was generated from another two *Ae. tauschii* accessions, Clae23 (**A**) and ENT336 **(B**). TPM values showed a strong correlation between the pairs of accessions. ($R^2$ = 0.7964 for Clae23 and 0.7932 for ENT336)

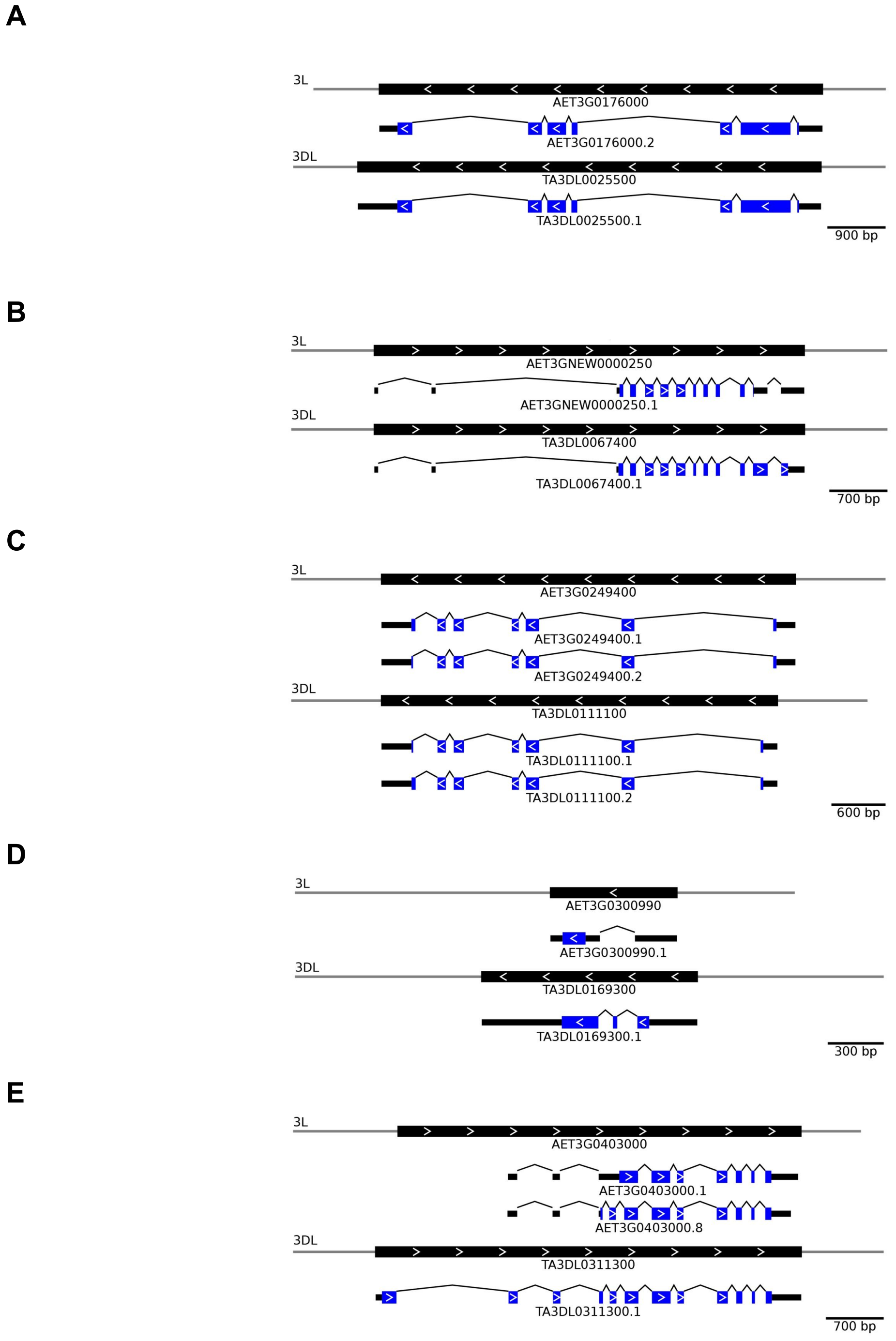


**Figure S3 Illustration of gene structure differences among 106 conserved DEGs between wheat and *Ae. tauschii.***

**A** An example of wheat TA3DL0025500 with the same structure as its syntenic gene pair in *Ae. tauschii*, and 4 gene pairs with different gene structures: B TA3DL0067400 (alternate start codon); **C** TA3DL0111100 (alternate first exon); **D** TA3DL0169300 (exon 2 in *Ae. tauschii* missing); and **E** TA3DL0311300 (*Ae. tauschii* has premature stop codon).

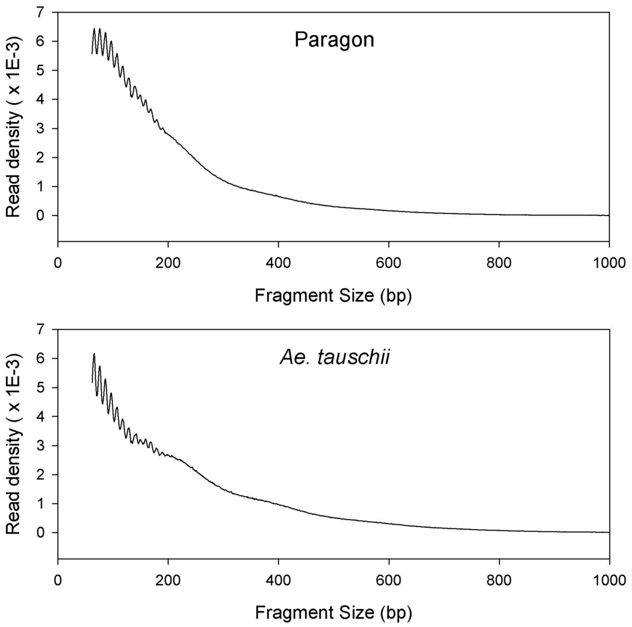


**Figure S4 Size distribution of ATAC-seq fragment lengths between *Triticum aestivum* Paragon and *Aegilops tauschii* AL8/78.**

The 10.5bp DNA pitch reflects the periodicity of right-handed helix in B-DNA. A trace of single and double nucleosome spacing can be seen in the 3L ATAC peaks, at approximately 200bp and 400bp.

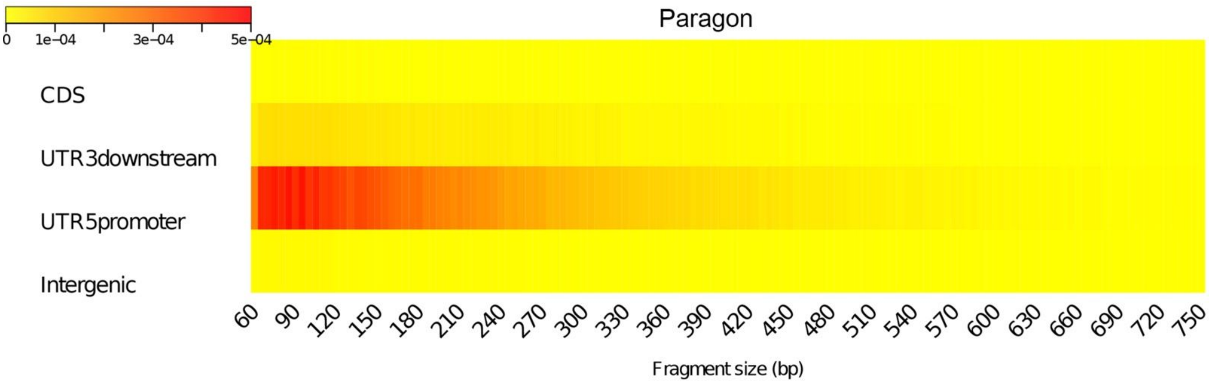


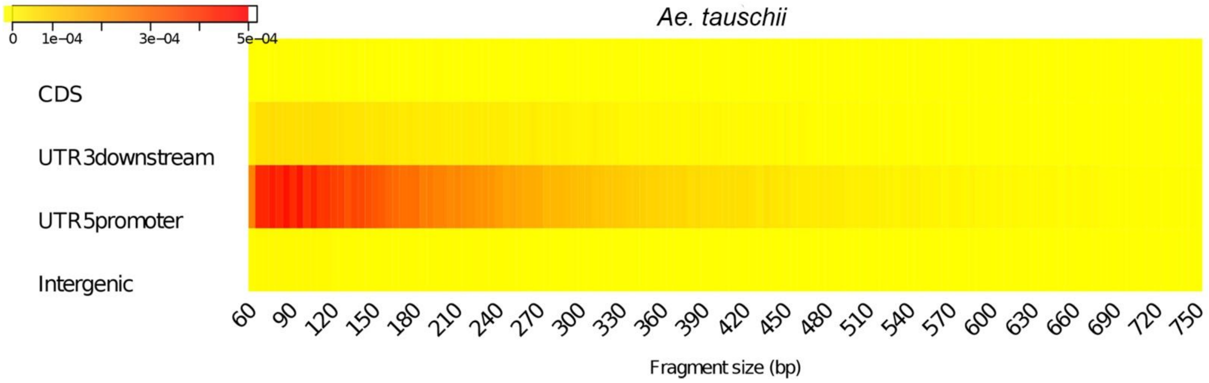


**Figure S5 Normalised read enrichments for four classes of chromosome states in *Triticum aestivum* Paragon and *Aegilops ta*uschii AL8/78.**

The normalised density plots of ATAC peaks showed that 5"UTR + promoter (2kb) regions had the highest read densities, followed by 3' UTR + downstream (2kb) regions, CDS + intron regions and intergenic regions.

**Table S1 Summary of repetitive elements of *Triticum aestivum* Chinese Spring 3DL and *Aegilops tauschii* AL8/78 3L**

| | | ***Triticum aestivum*** **chr3DL** | | ***Aegilops tauschii*** **chr3L** | |
|---|---|---|---|---|---|
| **Class** | **Subclass** | **Length (bp)** | **Percentage** | **Length (bp)** | **Percentage** |
| **DNA transposon** | | 45,694,659 | 12.29% | 47,299,305 | 12.54% |
| | CMC-ENSPM | 41,305,883 | 11.11% | 42,925,528 | 11.38% |
| **LINE** | | 1,670,347 | 0.45% | 1,651,400 | 0.44% |
| **LTR retrotransposon** | | 191,094,703 | 51.40% | 198,840,503 | 52.70% |
| | LTR-Copia | 56,109,154 | 15.09% | 57,910,706 | 15.35% |
| | LTR-Gypsy | 103,525,917 | 27.85% | 108,703,865 | 28.81% |
| **MobileElement** | | 956 | 0.00% | 847 | 0.00% |
| **RC** | | 58,706 | 0.02% | 73,186 | 0.02% |
| **rRNA** | | 19,149 | 0.01% | 17,408 | 0.00% |
| **Simple_repeat** | | 1,493,362 | 0.40% | 1,599,416 | 0.42% |
| **Low_complexity** | | 336,380 | 0.09% | 345,552 | 0.09% |
| **Other** | | 20,280,094 | 5.46% | 21,132,447 | 5.60% |
| **SUM** | | **257806071** | **69.35%** | **267937996** | **71.01%** |

**Table S2 Number of RNAseq read pairs after trimming in 5 sampled tissues of *Triticum aestivum* Paragon and *Aegilops tauschii* AL8/78**

| Tissues | Replicates | Paragon | AL8/78 |
|---|---|---|---|
| Leaves | Replicate 1 | 56,816,727 | 50,677,603 |
| | Replicate 2 | 56,452,015 | 55,377,053 |
| | Replicate 3 | 38,670,428 | 57,634,414 |
| Roots | Replicate 1 | 38,735,310 | 52,677,010 |
| | Replicate 2 | 51,869,501 | 55,016,084 |
| | Replicate 3 | 50,896,607 | 53,365,111 |
| Grains (10dd) | Replicate 1 | 59,835,166 | 80,624,534 |
| | Replicate 2 | 66,214,312 | 77,159,911 |
| | Replicate 3 | 57,335,248 | 68,751,597 |
| Grains (27dd) | Replicate 1 | 55,984,207 | 88,885,636 |
| | Replicate 2 | 51,331,203 | 75,465,851 |
| | Replicate 3 | 74,165,822 | 80,256,130 |
| Grains (Pooled) | Replicate 1 | 63,778,882 | 60,647,054 |
| Seedling (4dd) | Replicate 1 | 63,782,697 | 74,531,986 |
| | Replicate 2 | 77,936,789 | 71,175,660 |
| | Replicate 3 | 75,481,500 | 57,043,451 |

**Table S3 Number of differential expressed genes in 5 tissues between *Triticum aestivum* Paragon and *Aegilops tauschii* AL8/78**

| | Expressed genes | DEGs | Pseudogenes |
|---|---|---|---|
| Developing grain (10dd) | 1,716 | 277 (up 147, down 130) | 44 |
| Developing grain (27dd) | 1,516 | 173 (up 86, down 87) | 38 |
| Leaf | 1,564 | 262 (up 112, down 150) | 42 |
| Root | 1,784 | 327 (up 176, down 151) | 44 |
| Seedling | 1,829 | 251 (up 144, down 107) | 47 |
| Sum | 2,375 | 674 | 66 |

**Table S4 Mapping statistics after alignment of bisulfite treated *Triticum aestivum* Paragon and *Ae. tauschii* AL8/78 samples**

| | | Paragon | *Ae.tauschii* |
|---|---|---|---|
| Number of genes mapped | | 2,810 | 3,997 |
| C sites mapped on genes (min 10X) | | 846,450 | 458,347 |
| C sites mapped on 3DL promoters (min 10X) | | 418,818 | 289,521 |
| C's methylated overall | | | |
| | CpG | 89.9% | 87.1% |
| | CHG | 59.4% | 53.4% |
| | CHH | 3.8% | 3.5% |
| C's methylated genes | | | |
| | CpG | 66.7% | 41.2% |
| | CHG | 13.2% | 18.9% |
| | CHH | 1.0% | 3.8% |
| % converted | | 98.7 | 98.6 |

**Table S5 Number of regions for methylation analysis in *Triticum aestivum* Paragon 3DL and *Aegilops tauschii* AL8/78 3L**

| | Paragon | AL8/78 | Comparable regions |
|---|---|---|---|
| Gene CpG | 2,378 | 2,664 | 1,533 |
| Gene CHG | 2,467 | 2,643 | 1,586 |
| Gene CHH | 2,706 | 2,932 | 1,913 |
| Promoter CpG | 1,703 | 2,308 | 891 |
| Promoter CHG | 1,791 | 2,229 | 901 |
| Promoter CHH | 2,174 | 2,703 | 1,353 |

**Table S6 Summary of ATAC peaks and covered genes in different chromosome states of Paragon wheat and *Ae. tauschii* AL8/78**

| | Number of Peaks (Genes covered*) | |
|---|---|---|
| | **Paragon** | **AL8/78** |
| 5’UTR+promoter | 1,266 (1,098 genes) | 1,739 (869 genes) |
| CDS+Intron | 53 (42 genes) | 171 (130 genes) |
| 3’UTR+downstream | 226 (211 genes) | 480 (350 genes) |
| Intergenic | 1,425 | 2,570 |
| Total peaks | 2,970 (1,187 genes) | 4,960 (936 genes) |

*All genes including non-syntenic genes

**Table S7 Number of syntenic genes covered by ATAC-seq peaks**

| | Paragon | AL8/78 |
|---|---|---|
| Overall gene pairs | 930 genes (159 DEGs) | |
| Total | 774 (128 DEGs) | 362(88 DEGs) |
| CDS | 25(1 DEGs) | 46(13 DEGs) |
| UTR3downstream | 157(27 DEGs) | 132(35 DEGs) |
| UTR5promoter | 727(125 DEGs) | 345(85 DEGs) |
| Gene pairs with differential peaks | 816 genes (133 DEGs) | |
| CDS | 69 (14 DEGs) | |
| UTR3downstream | 251 (52 DEGs) | |
| UTR5promoter | 684 (98 DEGs) | |

**Table S8 Protein sequences used to evidence the exons in wheat chromosome 3DL annotation**

| Species | Version | Number | Sources |
|---|---|---|---|
| *Aegilops tauschii* | GCA_000347335.1 | 33,928 | Ensembl-25 |
| *Arabidopsis thaliana* | TAIR10 | 35,386 | Ensembl-25 |
| *Brachypodium distachyon* | v1.0 | 31,029 | Ensembl-25 |
| *Oryza sativa* | IRGSP-1.0 | 42,132 | Ensembl-25 |
| *Sorghium bicolor* | Sorbi1 | 36,338 | Ensembl-25 |
| *Triticum urartu* | GCA_000347455.1 | 33,483 | Ensembl-25 |
| *Hordeum vulgare* | 082214v1 | 62,311 | Ensembl-25 |
| *Triticum aesticum* | IWGSC2 | 99,354 | Ensembl-25 |
| *Triticum aesticum* | TriFLDB | 43,150 | http://trifldb.psc.riken.jp/v3 |
| *Zea mays* | AGPv3 | 63,235 | Ensembl-25 |
